# Assessing the validity of a calcifying oral biofilm model as a suitable proxy for dental calculus

**DOI:** 10.1101/2023.05.23.541904

**Authors:** Bjørn Peare Bartholdy, Irina M. Velsko, Shira Gur-Arieh, Zandra Fagernäs, Christina Warinner, Amanda G. Henry

## Abstract

Dental calculus is increasingly used by researchers to study dietary patterns in past populations. The benefits of using dental calculus for this purpose have been clearly demonstrated in previous studies, with dental calculus harbouring a wealth of microremains and biomarkers for health and diet within its mineral matrix. Previous studies have demonstrated some of the limitations and biases of how methods of processing may overlook, or even remove, some of the important information contained within the mineralised matrix. However, there are many factors that are impossible to account for *in vivo* and in archaeological material, such as exact dietary intake, and individual factors such as pH and enzyme activity, leaving some limitations that may not be addressed through these types of studies and will require a different approach.

We present a protocol for creating a calcifying oral biofilm model that can be used to explore the biases and limitations of dental calculus as a medium for paleodietary reconstructions. We report the microbial and mineral composition of our model in an effort to validate the model calculus as an appropriate proxy to natural dental calculus. The microbial profile and species diversity of our model was determined using metagenomic classification with the nf-core/eager pipeline and Kraken2, and compared to various reference samples from oral sites, including saliva, plaque, and dental calculus. We then assessed whether our model calculus mineralises in a manner similar to natural dental calculus using Fourier transform infrared (FTIR) spectroscopy. The metagenomic classification showed a microbial profile predominantly made up of (facultative) anaerobes, with a community structure that was somewhat distinct from other oral reference samples. The core genera of the model consisted of oral species, but clustered separately from oral reference samples, with a higher abundance of anaerobes.

Mineral and organic components of our model mimic that of the modern and archaeological reference calculus that was used as a comparison. There was an overall increase in the inorganic component relative to organic over the course of the experiment, with carbonated hydroxyapatite as the principal compound, consistent with natural human-derived calculus.

We conclude that oral biofilm models, such as the one presented in this study, have great potential to validate current methods used in the analysis of archaeological dental calculus, and should be used to complement, rather than replace current *in vivo* studies.

## 1 Introduction

Dental calculus is becoming an increasingly popular substance for exploring health and diet in past populations (Warinner et al., 2015). During life, dental plaque undergoes periodic mineralisation, trapping biomolecules and microfossils that are embedded within the dental plaque biofilm in the newly-formed dental calculus. This process is repeated as new plaque is deposited and subsequently mineralises, resulting in a layered structure representing a temporal record of biofilm growth and development (Warinner et al., 2014). The calculus serves as a protective casing for the entrapped biomolecules and microfossils, preserving them for thousands of years after death and burial (Fellows Yates et al., 2021). Studies using archaeological dental calculus span a wide range of topics in different regions and time periods. These include characterisation of the oral microbiome and its evolution in past populations (Adler et al., 2013; Fellows Yates et al., 2021; Kazarina et al., 2021; Velsko et al., 2019; Warinner et al., 2014), as well as extraction of microbotanical remains (Hardy et al., 2009; Henry & Piperno, 2008; Ma et al., 2022; Mickleburgh & Pagán-Jiménez, 2012) and other residues to infer dietary patterns and nicotine use (Bartholdy et al., 2023; Buckley et al., 2014; Eerkens et al., 2018; Hendy et al., 2018; Velsko, Overmyer, et al., 2017). Dental calculus has already provided a unique and valuable insight into the past, but the exact mechanism of the incorporation, retention, and preservation of microfossils and biomolecules exogenous to the microbial biofilm is largely unknown; even the process of plaque mineralisation is not fully understood (Jin & Yip, 2002; Omelon et al., 2013). This means that there may be hidden biases affecting our interpretations of dietary/activity patterns extrapolated from ancient dental calculus. These biases have been explored archaeologically (Fagernäs et al., 2022; Tromp et al., 2017) as well as in contemporary humans (Leonard et al., 2015) and non-human primates (Power et al., 2015), but not experimentally.

Dental plaque is an oral biofilm and is part of the normal state of the oral cavity. However, when left unchecked, plaque can lead to infections, such as dental caries and periodontitis, and/or mineralisation (Marsh, 2006). The dental plaque biofilm grows in a well-characterized manner before mineralisation, in a process that repeats regularly to build up dental calculus. Shortly after teeth are cleaned (whether mechanically or otherwise), salivary components adsorb to the crown or root and form the acquired dental pellicle. The pellicle provides a viable surface for bacteria to attach, especially early-coloniser species within the genera *Streptococcus* and *Actinomyces* (Marsh, 2006). Once the tooth surface has been populated by specialists in surface-attachment, other species of bacteria can attach to the adherent cells, increasing the biofilm density and diversity. The bacterial species secrete polysaccharides, proteins, lipids, and nucleic acids, into their immediate environment to form a matrix that provides structural support, nutrition, and allows for environmental niche partitioning (Flemming et al., 2016).

Biofilms can become susceptible to calcification under certain microenvironmental conditions, including an increased concentration of salts and a decrease in statherin and proline-rich proteins in saliva, rises in local plaque pH, and increased hydrolysis of urea (White, 1997; Wong et al., 2002). These conditions can cause increased precipitation and decreased dissolution of calcium phosphate salts within saliva and the plaque biofilm. The resulting supersaturation of calcium phosphate salts is the main driver of biofilm mineralisation (Jin & Yip, 2002). The primary minerals in dental calculus are hydroxyapatite, octacalcium phosphate, whitlockite, and brushite. During initial mineralisation the main mineral component is brushite, which shifts to hydroxyapatite in more mature dental calculus (Hayashizaki et al., 2008; Jin & Yip, 2002). The exact elemental composition of dental calculus varies among individuals due to various factors, including diet (Hayashizaki et al., 2008; Ji et al., 2000).

Dental plaque can also be grown *in vitro*, and these oral biofilm models are commonly used in dental research to assess the efficacy of certain treatments on dental pathogens (Exterkate et al., 2010; Filoche et al., 2007) without the ethical issues of inducing plaque accumulation in study participants and the complexity of access and sampling in humans or animals. Oral biofilm models are often short-term models grown over a few days, but longer term models also exist (up to six weeks) which are used to develop mature plaque or dental calculus (Middleton, 1965; Sissons et al., 1991; Velsko & Shaddox, 2018; Wong et al., 2002). A well-known limitation of biofilm models is the difficulty in capturing the diversity and complexity of bacterial communities and metabolic dependencies, micro-environments, nutrient availability, and host immune-responses in the natural oral biome (Bjarnsholt et al., 2013; Edlund et al., 2018; Velsko, Cruz-Almeida, et al., 2017; Velsko & Shaddox, 2018). These limitations can be overcome by complex experimental setups, but at the cost of lower throughput and increased requirements for laboratory facilities.

Despite the limitations, oral biofilm models have many benefits over *in situ* research. There are many variables involved in dental calculus formation, such as intra- and inter-individual variation in salivary flow, oral pH, and amylase activity, which can be hard to tease apart *in situ*. Oral biofilm models provide a controlled environment to explore the effect of selected variables on the growth of calculus and the retention of dietary components in the biofilm, as well as a means to identify how the methods used in archaeology may inadvertently bias the interpretations. This type of research has, so far, been limited, but has the potential to greatly benefit archaeological research on past diet (Radini & Nikita, 2022).

We present an oral biofilm model that can serve as a viable proxy for dental calculus for archaeology-oriented research questions. It is a multispecies biofilm using whole saliva as the inoculate, with a simple multiwell plate setup that is accessible even to smaller lab budgets and those with limited facilities for microbiology work. Here, we used next-generation sequencing and metagenomic classification to characterise the bacterial composition of our model dental calculus and compare it to oral reference samples, including saliva, buccal mucosa, plaque, and modern human dental calculus. This was done to ensure that the model microbiome is predominantly oral and not overgrown by environmental contaminants. We then determined the mineral composition of the model dental calculus using Fourier transform infrared (FTIR) spectroscopy to verify the presence of calculus-specific mineral phases and functional groups, and perform a qualitative comparison with modern and archaeological reference calculus. Overall the model calculus is chemically similar to natural calculus, and has a predominantly oral microbiome. The microbial diversity and richness within the model samples were lower than oral reference samples, suggesting that the model samples do not contain identical species composition and abundances as the natural samples. The mineral composition closely resembles modern and archaeological reference calculus, predominantly comprised of carbonate hydroxyapatite with a similar level of crystallinity and order. As such, the model dental calculus presented here is a viable proxy to natural dental calculus and can be used to explore many of the currently unexplained processes we see in the archaeological material, when working within the limitations of an oral biofilm model.

## 2 Materials and Methods

Our biofilm setup consists of whole saliva as the inoculate to approximate natural microbial communities within the human oral cavity, and a 24-well plate to generate multiple replicated conditions in a single experimental run (see Figure 1 for an overview of the protocol). The biofilm is grown for 25 days to allow time for growth of larger deposits and mineralisation. Raw potato and wheat starch solutions were added during the biofilm growth to explore the biases involved in their incorporation and extraction from dental calculus. These results are presented in a separate article (Bartholdy & Henry, 2022).

**Figure 1:**
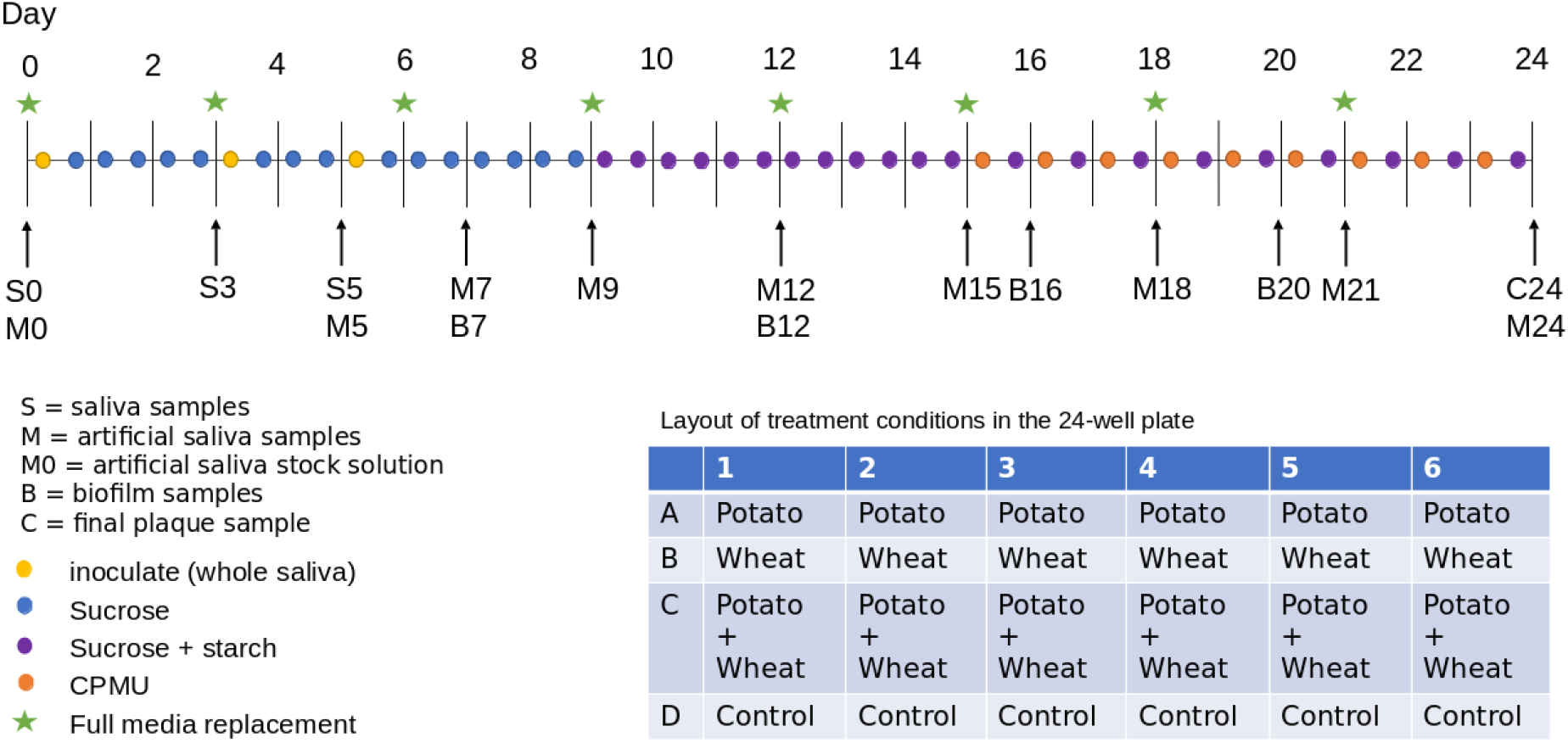
Overview of the protocol for biofilm growth. The samples for metagenomic analysis were grown in a separate experimental plate than the FTIR samples under the same experimental conditions. Biofilm (B) and calculus (C) samples were used for FTIR spectroscopy, and saliva (S), artificial saliva (M), and calculus samples were used for metagenomic analysis.

To determine the composition of microbial communities, we sampled the medium from the biofilm wells over the course of the experiment. We sequenced the DNA to identify species that are present in the model, and assess whether these mimic natural oral communities. During a separate experimental run, under the same conditions, we directly sampled the biofilms on multiple days and determined the mineral composition using FTIR, and compared the spectra to those of natural dental calculus, both modern and archaeological. Samples were taken from both controls and starch treatments, but differences between these samples were not explored in this study.

### 2.1 Biofilm growth

We employ a multispecies oral biofilm model following a modified protocol from Sissons and colleagues (1991) and Shellis (1978). The setup comprises a polypropylene 24 deepwell PCR plate (KingFisher 97003510) with a lid containing 24 pegs (substrata), which are autoclaved at 120°C, 1 bar overpressure, for 20 mins.

The artificial saliva (hereafter referred to as medium) is a modified version of the basal medium mucin (BMM) described by Sissons and colleagues (1991). It is a complex medium containing 2.5 g/l partially purified mucin from porcine stomach (Type III, Sigma M1778), 5 g/l trypticase peptone (Roth 2363.1), 10 g/l proteose peptone (Oxoid LP0085), 5 g/l yeast extract (BD 211921), 2.5 g/l KCl, 0.35 g/l NaCl, 1.8 mmol/l CaCl_2_, 5.2 mmol/l Na_2_HPO_4_ (Sissons et al., 1991), 6.4 mmol/l NaHCO_3_ (Shellis, 1978), 2.5 mg/l haemin. This is subsequently adjusted to pH 7 with NaOH pellets and stirring, autoclaved (15 min, 120°C, 1 bar overpressure), and supplemented with 5.8 (mu)mol/l menadione, 5 mmol/l urea, and 1 mmol/l arginine (Sissons et al., 1991).

Fresh whole saliva (WS) for inoculation was provided by a 31-year-old male donor with no history of caries, who abstained from oral hygiene for 24 hours, and no food was consumed two hours prior to donation. No antibiotics were taken up to six months prior to donation. Saliva was stimulated by chewing on parafilm, then filtered through a bleach-sterilised nylon cloth to remove particulates. Substrata were inoculated with 1 ml/well of a two-fold dilution of WS in sterilised 20% glycerine for four hours at 36°C, to allow attachment of the salivary pellicle and plaque-forming bacteria. After initial inoculation, the substrata were transferred to a new plate containing 1 ml/well medium and incubated at 36°C, with gentle motion at 30 rpm. The inoculation process was repeated on days 3 and 5 by transferring the samples to a new plate with inoculate. Medium was partially refreshed once per day, by topping up the wells to the original volume with more medium, and fully refreshed every three days, throughout the experiment, by transferring the substrata to a new plate containing medium. To feed the bacteria, the substrata were transferred to a new plate, containing 5% (w/v) sucrose, for six minutes twice daily, except on inoculation days (days 0, 3, and 5), where the samples only received one sucrose treatment after inoculation.

On day 9, starch treatments were introduced, replacing sucrose treatments (except for control sample). As with the sucrose treatments, starch treatments occurred twice per day for six minutes, and involved transferring the substrata to a new plate containing a 0.25% (w/v) starch from potato (Roth 9441.1) solution, a 0.25% (w/v) starch from wheat (Sigma S5127) solution, and a 0.5% (w/v) mixture of equal concentrations (w/v) wheat and potato. All starch solutions were created in a 5% (w/v) sucrose solution. Before transferring biofilm samples to the starch treatments, the starch plates were agitated to keep the starches in suspension in the solutions, and during treatments, the rpm was increased to 60. The purpose of starch treatments was to explore the incorporation of starch granules into the model calculus. Starch treatments were initiated on day 9 (Figure 1) to avoid starch granule counts being affected by *α*-amylase hydrolysis from the inoculation saliva. An *α*-amylase assay conducted on samples from days 3, 6, 8, 9, 10, 12, and 14 also showed that there was no host salivary *α*-amylase activity in the system. The results of the starch incorporation and *α*-amylase activity assay have been reported in a separate article (Bartholdy & Henry, 2022).

After 15 days, mineralisation was encouraged with a calcium phosphate monofluorophosphate urea (CPMU) solution containing 20 mmol/l CaCl_2_, 12 mmol/l NaH_2_PO_4_, 5 mmol/l Na_2_PO_3_F, 500 mmol/l urea (Pearce & Sissons, 1987; Sissons et al., 1991), and 0.04 g/l MgCl. The substrata were submerged in 1 ml/well CPMU five times daily, every two hours, for six minutes, at 30 rpm. During the mineralisation period, starch treatments were reduced to once per day, two hours after the last CPMU treatment. This cycle was repeated for 10 days until the end of the experiment on day 24 (Figure 1). More detailed protocols are available at https://dx.doi.org/10.17504/protocols.io.dm6gpj9rdgzp/v1.

All laboratory work was conducted in sterile conditions under a laminar flow hood to prevent starch and bacterial contamination. Starch-free control samples that were only fed sucrose were included to detect starch contamination.

### 2.2 Metagenomics

A total of 35 samples were taken during the experiment from the donated saliva, artificial saliva, and from the biofilm end-product on day 24 (Table 1). DNA extraction was performed at the Max Planck Institute for the Science of Human History (Jena, Germany), using the DNeasy PowerSoil Kit from QIAGEN. C2 inhibitor removal step skipped, going directly to C3 step.

**Table 1:** Number of samples taken during the experiment, separated by sampling day and sample type.

| Sample type | Sampling day | n |
| --- | --- | --- |
| saliva | 0 | 1 |
| saliva | 3 | 1 |
| saliva | 5 | 1 |
| medium | 5 | 2 |
| medium | 7 | 2 |
| medium | 9 | 2 |
| medium | 12 | 2 |
| medium | 15 | 2 |
| medium | 18 | 2 |
| medium | 21 | 2 |
| medium | 24 | 2 |
| model_calculus | 24 | 16 |

The DNA was sheared to 500bp through sonication with a Covaris M220 Focused-ultrasonicator. Double-stranded libraries were prepared (Aron et al., 2020) and dual indexed (Stahl et al., 2019), with the indexing protocol being adapted for longer DNA fragments. Briefly, the modifications consisted of adding 3 l of DMSO to the indexing reaction, and extending the amplification cycles to 95°C for 60 s, 58°C for 60 s, and 72°C for 90 s. The libraries were paired-end sequenced on a NextSeq 500 to 150bp, and demultiplexed by an in-house script.

#### 2.2.1 Preprocessing

The raw DNA reads were preprocessed using the nf-core/eager, v2.4.4 pipeline (Fellows Yates et al., 2020). The pipeline included adapter removal and read merging using AdapterRemoval, v2.3.2 (Schubert et al., 2016). Merged reads were mapped to the human reference genome (GRCh38) using BWA, v0.7.17-r1188 (Li & Durbin, 2009) (-n 0.01; -l 32), and unmapped reads were extracted using Samtools, v1.12. The final step of the pipeline, metagenomic classification, was conducted in kraken, v2.1.2 (Wood et al., 2019) using the Standard 60GB database (https://genome-idx.s3.amazonaws.com/kraken/k2_standard_20220926.tar.gz).

Environmental reference samples were downloaded directly from ENA and from NCBI using the SRA Toolkit. Oral reference samples were downloaded from the Human Metagenome Project (HMP), and modern calculus samples from Velsko et al. (2017). From the HMP data, only paired reads were processed, singletons were removed. *In vitro* biofilm model samples from Edlund et al. (2018) were used as a reference. Links to the specific sequences are included in the metadata. Human-filtered reads produced in this study were uploaded to ENA under accession number PRJEB61886.

#### 2.2.2 Authentication

Species with lower than 0.001% relative abundance across all samples were removed from the species table. Source-Tracker2 (Knights et al., 2011) was used to estimate source composition of the abundance-filtered oral biofilm model samples using a Bayesian framework, and samples falling below 70% oral source were removed from downstream analyses. Well-preserved abundance-filtered samples were compared to oral and environmental controls to detect potential external contamination. The R package decontam v1.20.0 (Davis et al., 2018) was used to identify potential contaminants in the abundance-filtered table using DNA concentrations with a probability threshold of 0.95 and negative controls with a probability threshold of 0.05. Putative contaminant species were filtered out of the OTU tables for all downstream analyses.

#### 2.2.3 Community composition

Relative abundances of communities were calculated at the species- and genus-level, as recommended for compositional data (Gloor et al., 2017). Shannon index and Pileou’s evenness index were calculated on species-level OTU tables of all model and oral reference samples using the vegan v2.6.4 R package (Oksanen et al., 2022). Shannon index was calculated for all experimental samples to see if there is an overall loss or gain in diversity and richness across the experiment. Sparse principal component analysis (sPCA) was performed on model biofilm samples to assess differences in microbial composition between samples within the experiment, and a separate sPCA analysis was performed on model calculus and oral reference samples. The sPCA analysis was conducted using the mixOmics v 6.24.0 R package (Rohart et al., 2017).

The core microbiome was calculated by taking the mean genus-level relative abundance within each sample type for model calculus, modern reference calculus, sub- and supragingival plaque. Genera present at lower than 5% relative abundance were grouped into the category ‘other’. Information on the oxygen tolerance of bacterial species was downloaded from BacDive (Reimer et al., 2022) and all variations of the major categories anaerobe, facultative anaerobe, and aerobe were combined into the appropriate major category. At the time of writing, 55.7% species were missing aerotolerance values. This was mitigated by aggregating genus-level tolerances to species with missing values, and may have some errors (although unlikely to make any significant difference).

#### 2.2.4 Differential abundance

Differential abundance of species was calculated using the Analysis of Compositions of Microbiomes with Bias Correction (ANCOM-BC) method from the ANCOMBC R package v2.2.0 (Lin & Peddada, 2020), with a species-level OTU table as input. Results are presented as the log fold change of species between paired sample types with 95% confidence intervals. P-values are adjusted using the false discovery rate (FDR) method. Samples are grouped by sample type (i.e. saliva, plaque, modern calculus, model calculus). To supplement the sPCA analyses, we visualised the log-fold change of the top 30 species in each of principal components 1 and 2, allowing us to see which species are enriched in the different samples and causing clustering in the sPCA.

### 2.3 FTIR

To determine the mineral composition and level of crystallisation of the model dental calculus samples, we used Fourier Transform Infrared (FTIR) spectroscopy. We compared the spectra of model dental calculus with spectra of archaeological and modern dental calculus and used a built-in Omnic search library for mineral identification (Mentzer et al., 2014; Weiner, 2010b). The archaeological dental calculus was sampled from an isolated permanent tooth from Middenbeemster, a rural, 19th century Dutch site (Lemmers et al., 2013). Samples were analysed at the Laboratory for Sedimentary Archaeology, Haifa University. The analysis was conducted with a Thermo Scientific Nicolet is5 spectrometer in transmission, at 4 cm^−1^ resolution, with an average of 32 scans between 4000 and 400 cm^−1^ wavenumbers.

Analysis was conducted on 28 model calculus samples from days 7, 12, 16, 20, and 24 (Table 2). Some samples from the same sampling day had to be combined to provide enough material for analysis. Samples analysed with FTIR were grown during a separate experimental run from the samples sequenced for DNA, but following the same setup and protocol (as described above). Samples were analysed following the method presented in Asscher, Regev, et al. (2011) and Asscher, Weiner, et al. (2011). A few *μ*g of each sample were repeatedly ground together with KBr and pressed in a 7 mm die under two tons of pressure using a Specac mini-pellet press (Specac Ltd., GS01152). Repeated measurements of the splitting factor (SF) of the absorbance bands at 605 and 567 cm1 wavenumbers were taken after each grind, and a grind curve was produced following Asscher, Regev, et al. (2011) to try and detect changes in the hydroxyapatite crystallinity over time. Samples were ground and analysed up to six times (sample suffix a-f) for the grinding curve. Grinding curves were prepared for samples from days 16, 20, and 24. No grind curves were produced for samples from days 7 and 12. These were largely composed of organics and proteins, and did not form enough mineral (hydroxyapatite) for analysis. The splitting factor of carbonate hydroxyapatite was calculated using a macro script, following Weiner & Bar-Yosef (1990). The calculation involves dividing the sum of the height of the absorptions at 603 cm^−1^ and 567 cm^−1^ by the height of the valley between them. Following Asscher, Regev, et al. (2011) and Asscher, Weiner, et al. (2011), we plotted the splitting factor against the full width at half maximum (FWHM) of the main absorption at 1035-1043 cm^−1^ to explore crystallinity (crystal size) and the order and disorder of hydroxyapatite. We then compared our grinding curve slopes and FWHM to the ones produced by Asscher, Weiner, et al. (2011). Asscher, Weiner, et al. (2011) and Asscher, Regev, et al. (2011) demonstrated that while the decrease in FWHM of each grinding in the curve reflects a decrease in particle size due to grinding, the location of the curves within a plot of the FWHM against the splitting factor expresses the disorder effect. Thus the curves with steeper slopes, higher splitting factor, and lower FWHM represent lower levels of disorder in the mineral (Figure 2 in Asscher, Weiner, et al., 2011).

**Table 2:** Summary of samples used in FTIR analysis, including type of sample, sampling day, number of samples (n), and mean weight in mg.

| Sample type | Sampling day | n | Weight (mg) |
| --- | --- | --- | --- |
| model_biofilm | 7 | 3 | 0.59 |
| model_biofilm | 12 | 4 | 1.09 |
| model_biofilm | 16 | 7 | 2.00 |
| model_biofilm | 20 | 6 | 3.50 |
| model_calculus | 24 | 8 | 3.87 |

**Figure 2:**
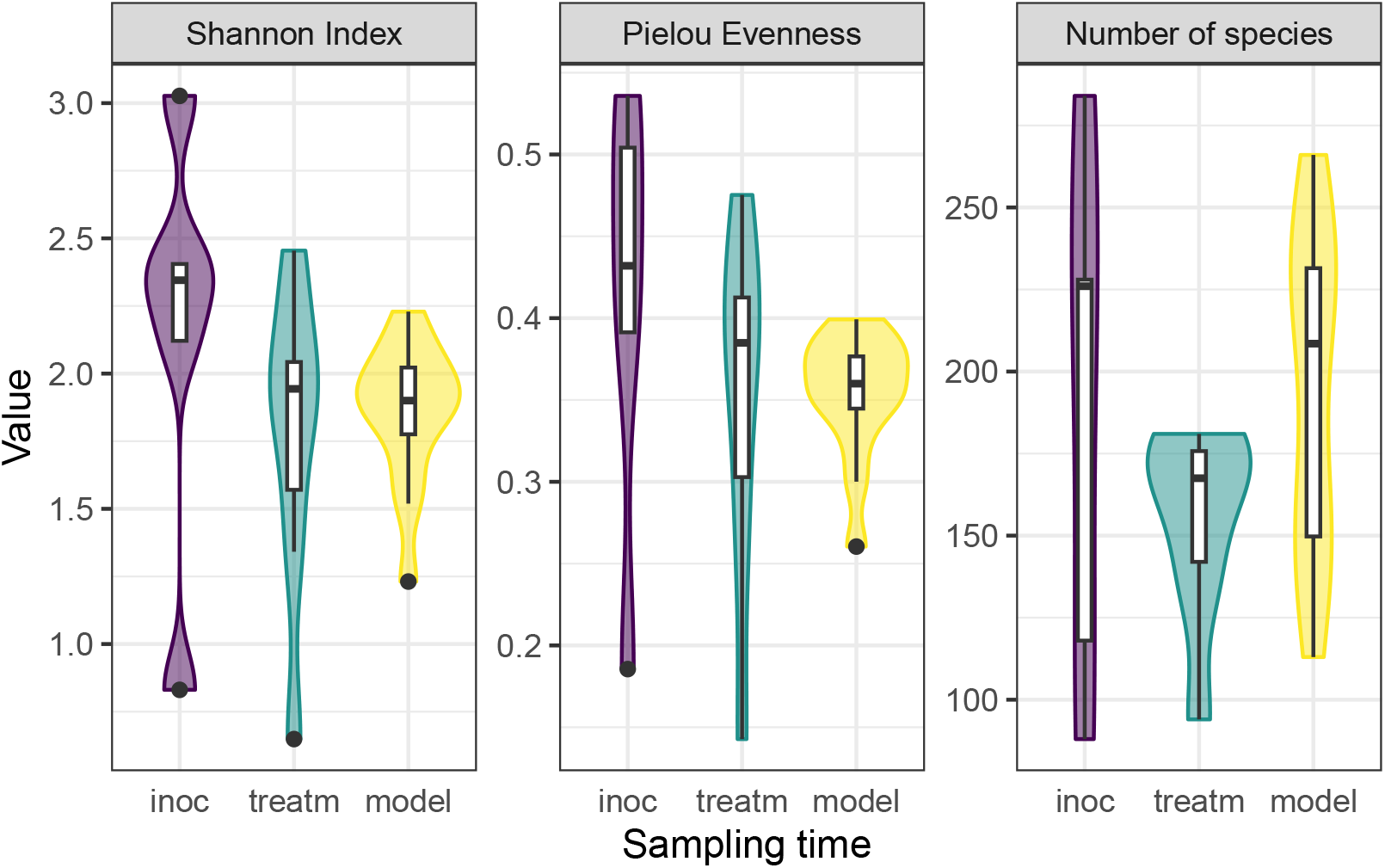
Plot of Pielou Evenness Index, number of species, and Shannon Index across experiment samples grouped by sampling time. inoc = samples from days 0-5; treatm = samples from days 6-23; model = model calculus samples from day 24.

### 2.4 Statistics

Statistical analysis was conducted in R version 4.3.0 (2023-04-21) (Already Tomorrow) (R Core Team, 2020). Data cleaning and wrangling performed with packages from tidyverse (Hadley Wickham et al., 2019). Plots were created using ggplot2 v3.4.2 (H. Wickham, 2016).

## 3 Results

### 3.1 Metagenomic analysis

#### 3.1.1 Sample authentication

To determine the extent of contamination in our samples, we performed a source-tracking analysis using Source-Tracker2 (Knights et al., 2011). Results suggest that the majority of taxa across samples have an oral microbial signature, and therefore our samples are minimally affected by external contamination (Figure S1). We compared SourceTracker2 results to a database of oral taxa from the cuperdec v1.1.0 R package (Fellows Yates et al., 2021) to prevent removal of samples where oral taxa were assigned to a non-oral source (Figure S2), as some taxa with a signature from multiple sources are often classified as “Unknown” (Velsko et al., 2019). We included several oral sources, which may increase the risk of this occurring. Samples containing a large proportion (>70%) of environmental contamination were removed. The removed samples were predominantly medium samples from later in the experiment, and a few model calculus samples. After contaminated samples were removed, suspected contaminant-species were removed from the remaining samples using the decontam R package (Davis et al., 2018). After contamination removal, samples consisted of between 88 and 284 species with a mean of 182.

#### 3.1.2 Decrease in community diversity across experiment

To monitor the development of microbial communities over the course of the experiment, we used the Shannon Index to assess the species diversity and richness at various stages of our protocol. Samples were grouped into sampling categories due to low sample sizes on sampling days (inoc = days 0, 3, 5; treatm = days 7, 9, 12, 15; model = day 24). There was a slight decrease in mean Shannon Index between inoculation and treatment samples, followed by a slight increase to model calculus samples, as well as a decrease in variance within samples types. The Pielou Evenness Index showed a similar pattern while the number of species increased between the treatment period and the final model calculus (Figure 2).

#### 3.1.3 Medium and model calculus samples are distinct from the inoculate

We next examined whether there is a change in the species composition over time in our samples by assessing the beta-diversity in a PCA. The species profiles of the saliva inoculate used in our experiment were distinct from both medium and model calculus samples. Most of the separation of saliva from model calculus is on PC1 of the sPCA, where most of the positive sample loadings are driven by anaerobic species (model calculus), especially *Selenomonas* spp, and negative loadings are predominantly facultative anaerobes and some aerobes, such as *Rothia* and *Neisseria* spp (saliva). Medium and saliva are separated mostly on PC2, with medium samples located between saliva and model calculus samples. Model calculus samples also cluster separately from the medium samples on PC2, with some overlap between the more mature medium samples and model calculus. Most of the negative loadings separating saliva and model calculus from medium samples are dominated by *Actinomyces* spp., while positive species loadings are more diverse, and seemingly unrelated to aerotolerance (Figure 3).

**Figure 3:**
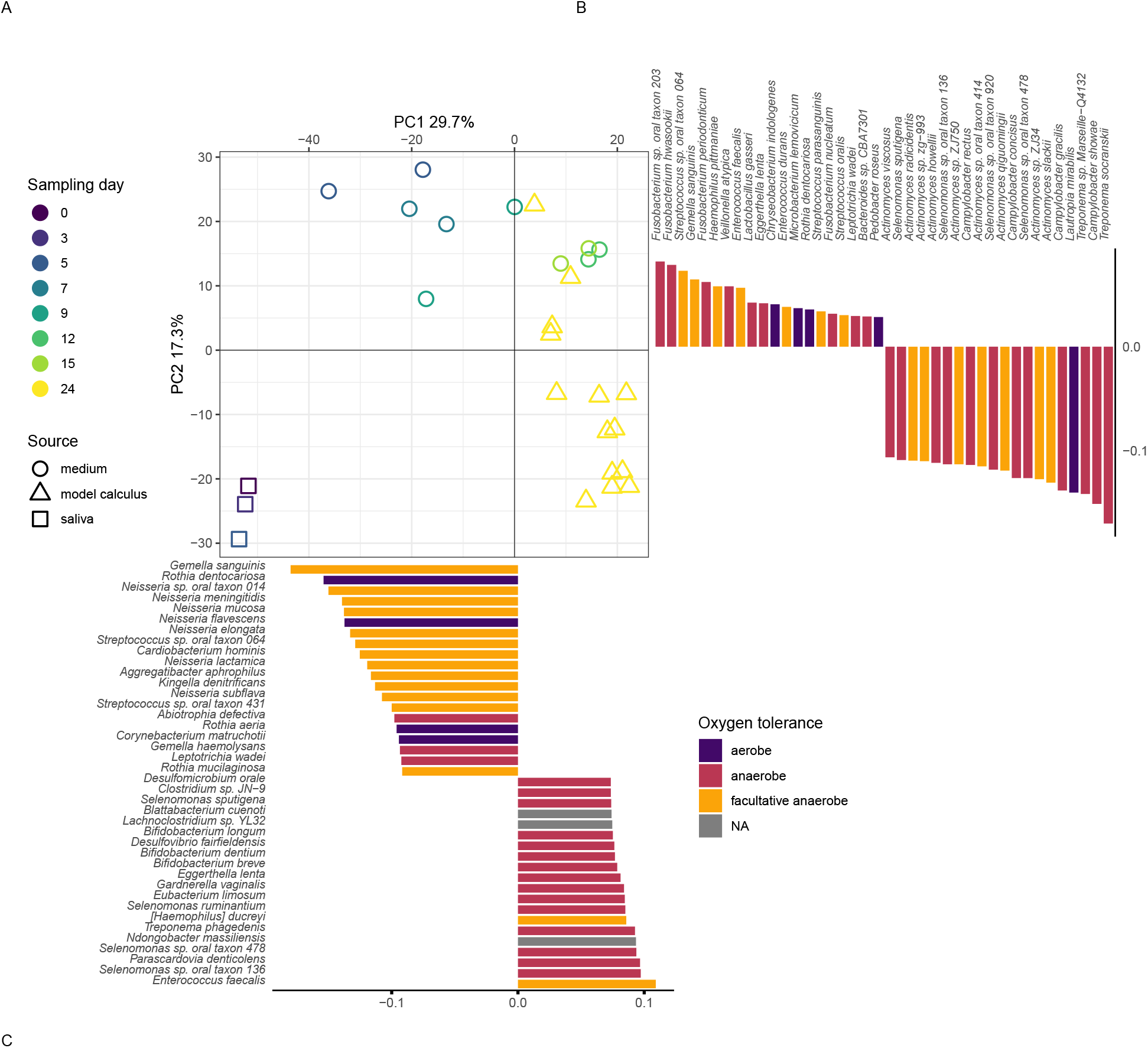
sPCA on species-level counts and oxygen tolerance in samples from this study only. Figure shows the main sPCA plot (A), species loadings on PC2 (B), and species loadings on PC1 (C).

We determined whether there are species that are differentially abundant between our sample types using the AN-COMBC R package (Lin & Peddada, 2020), giving us an idea of how the biofilm develops under our experimental conditions. Species enriched in saliva compared to model calculus are largely aerobic or facultatively anaerobic, while species enriched in model calculus compared to saliva are mainly anaerobes. The differences between saliva and calculus are more pronounced than between medium and model calculus, which is expected (Figure 4).

**Figure 4:**
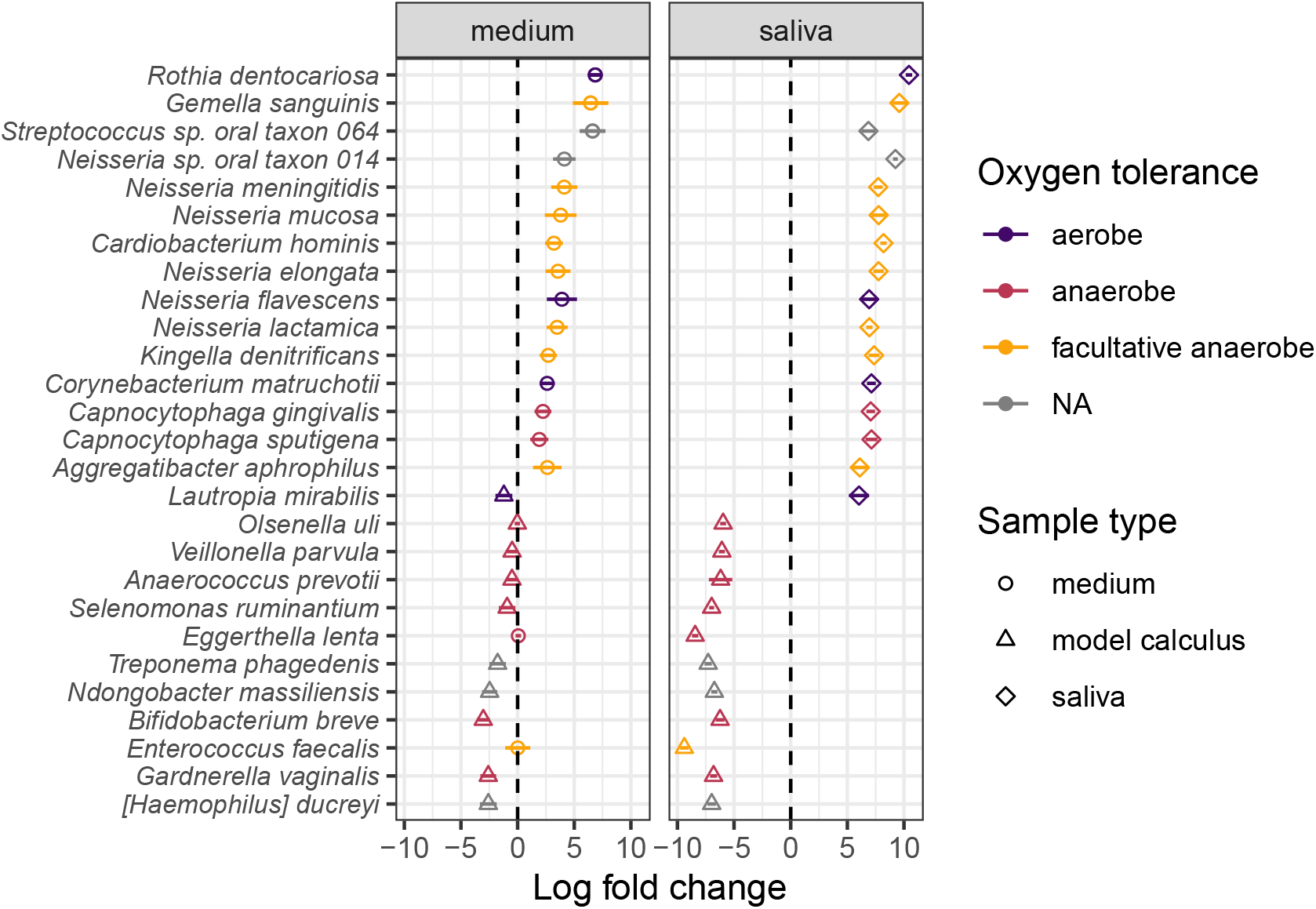
Log-fold changes between sample types. Circles are species enriched in the model calculus, squares are enriched in saliva, and triangles in medium. Lines are standard error. Plot shows the top 30 absolute log-fold changes between model calculus and saliva.

#### 3.1.4 Lower diversity in artificial samples than oral references

We used the Shannon Index to compare alpha-diversity in our model to oral reference samples. The mean Shannon Index of model samples—medium, model calculus, reference *in vitro* biofilm were consistently lower than the means of oral reference samples—mucosa, modern reference dental calculus, saliva, and subgingival and subgingival plaque. The Pielou species evenness index has a similar distribution, although the comparative biofilm samples have a higher mean than biofilm samples from this study. Saliva inoculate samples from this study have a lower mean Shannon index than reference samples, which may have contributed to the lower alpha-diversity in model samples compared to reference samples. The number of species follows the same trend.

#### 3.1.5 Model calculus is distinct from dental calculus and other oral samples

We calculated the mean relative abundances of the genera in each sample to compare the core genera of model calculus with oral reference samples. The most common genera (>5% relative abundance) are shown in Figure 6. The main overlap between the model calculus and oral reference samples is the high relative abundance of *Streptococcus*. Model calculus consists mostly of *Enterococcus* and *Veillonella* spp., despite both having low abundance in donor saliva. *Enterococcus* are also known environmental contaminants, and we cannot exclude environmental contamination as a possible source for these species in our model. Oral reference samples have a more balanced composition, as they are also represented by fastidious early-coloniser species like *Capnocytophaga* and *Neisseria* spp., which require an environment with at least 5% carbon dioxide to thrive (Tønjum & van Putten, 2017).

**Figure 5:**
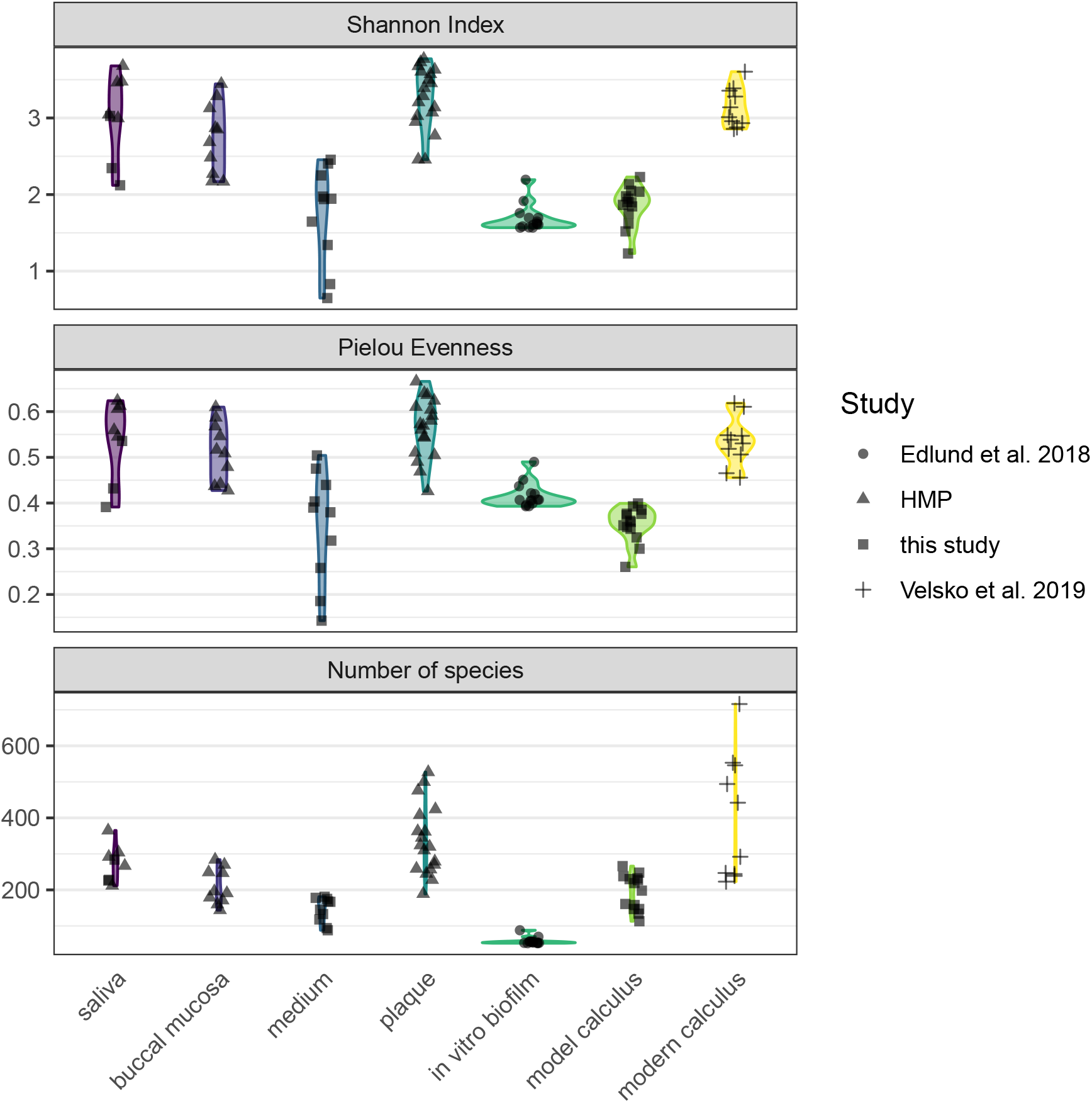
Shannon Index for model calculus and medium samples, as well as oral reference samples and comparative *in vitro* study.

**Figure 6:**
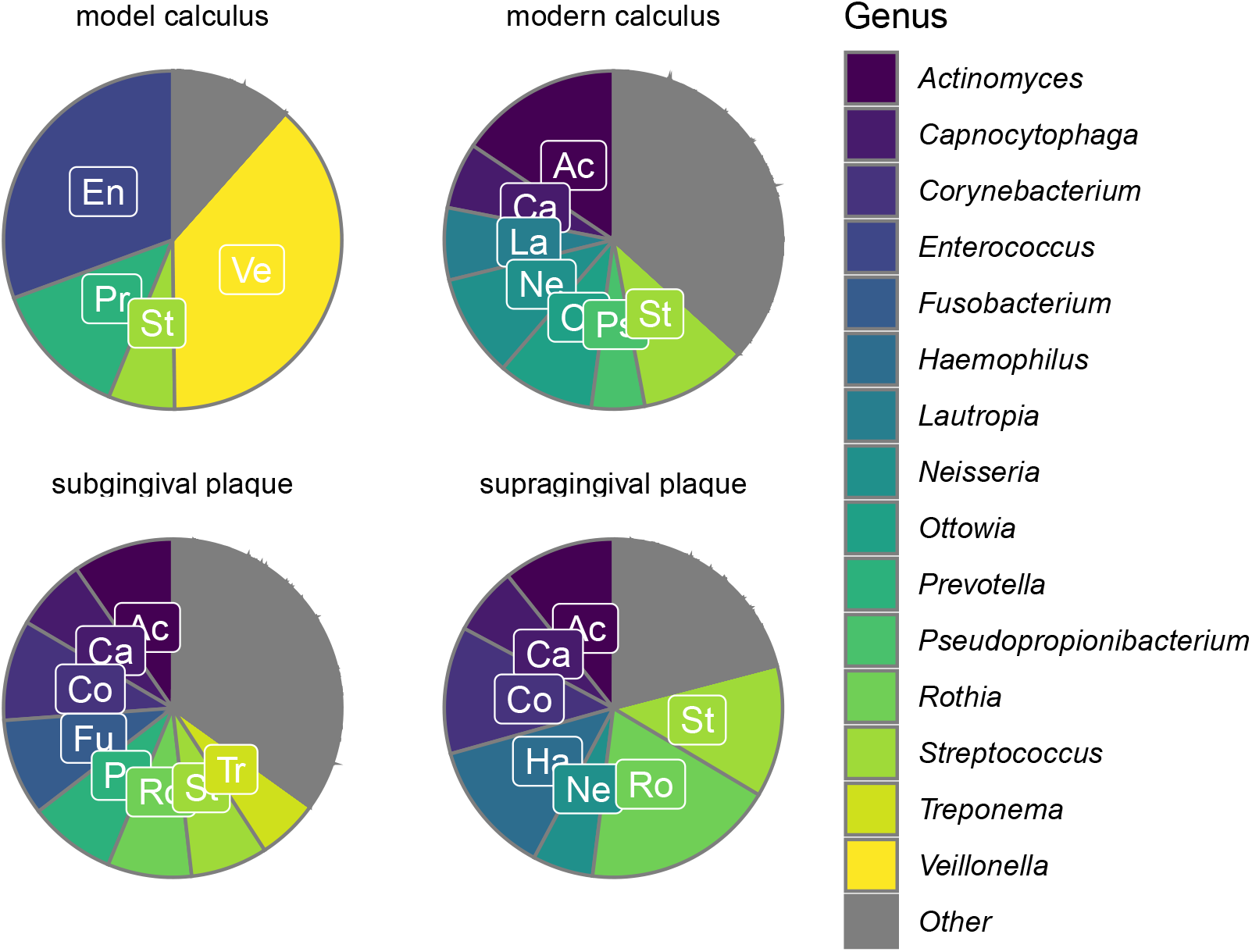
Core genera within the different types of samples represented as mean relative abundances at the genus level. Other = other genera present in lower than 5% relative abundance.

To directly compare the beta-diversity of our model calculus with oral reference samples, including modern dental calculus, we used an sPCA including only our model calculus and reference samples. Model calculus samples are distinct from both the oral reference samples and the biofilm model reference samples. They are separated from oral reference samples mainly on PC1, and from biofilm model reference samples (and, to some extent, oral samples) on PC2. The highest negative contributions are a mix of all types of aerotolerance, while the positive contributions are mostly (facultative) anaerobes, with *Enterococcus* spp. as the top three positive contributors to PC1. Top negative contributors are *Capnocytophaga* spp as well as the aerobes *Corynebacterium matruchotii* and *Rothia dentocariosa*. The top positive contributors to PC2 are all anaerobes, mainly from the genus *Selenomonas*. Top negative contributors to PC2 are a mix of aerotolerances, with many *Streptococcus* spp.

To investigate which species are enriched in different sample types, and compare the final product of our model with naturally occurring plaque and calculus samples, we performed differential abundance analysis on our model calculus samples, modern dental calculus, and sub- and supragingival plaque. Based on the differential abundance analysis the main differences between model calculus and oral reference samples, when looking at the top 30 contributors to PC1, are that the oral reference samples are enriched with species with a diverse oxygen tolerance from a wide range of genera, while the model calculus is enriched with *Enterococcus* spp. The largest differences occur in *Corynebacterium matruchotii, Rothia dentocariosa*, and *Capnocytophaga gingivalis* (Figure 8A). This is echoed when looking at the top 30 contributors to PC2, where most of the species are enriched in model calculus, all of which are anaerobes, and the largest differences occurring in *Cryptobacterium curtum, Eggerthella lenta*, and *Mogibacterium diversum* (Figure 8B).

**Figure 7:**
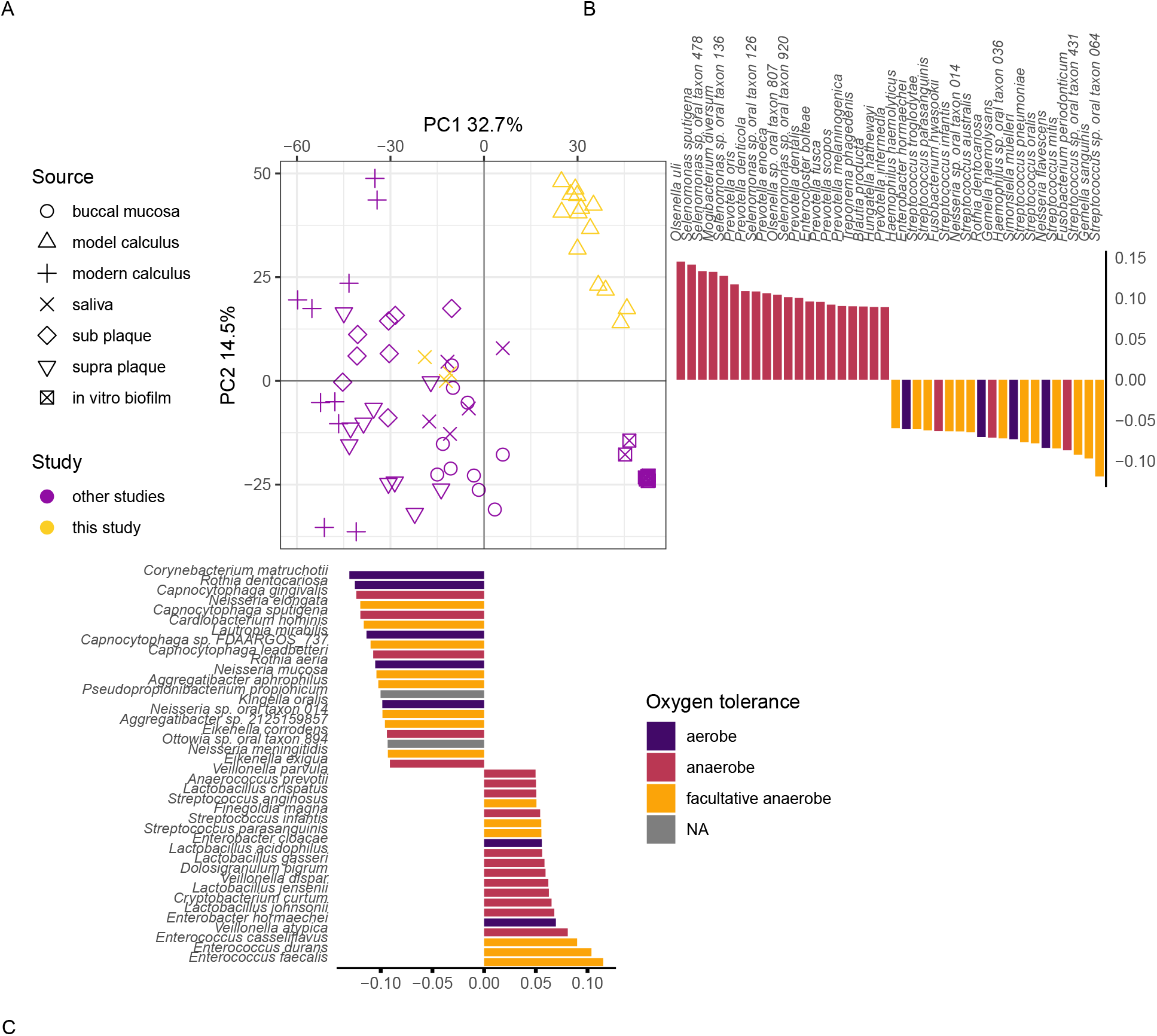
sPCA on species-level counts from model calculus and reference samples. Figure shows (A) the main sPCA plot, (B) the species loadings from PC2, and (C) species loadings on PC1.

**Figure 8:**
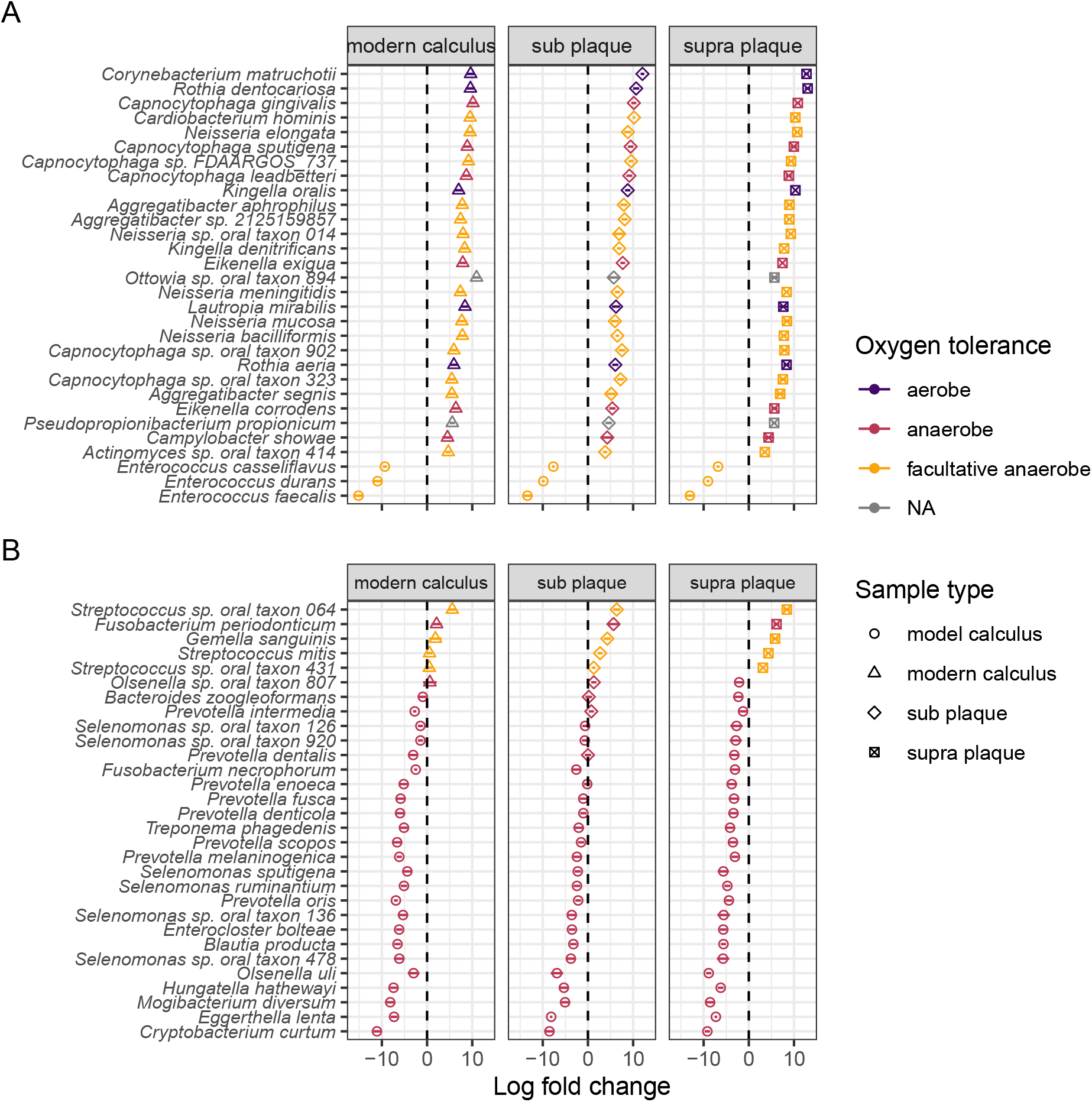
Log-fold changes between sample types. Circles are species enriched in the model calculus, triangles in modern calculus, diamonds are enriched in subgingival plaque, and squares in supragingival plaque. Plot shows the top 30 loadings (absolute value) in PC1 (A) and PC2 (B) between model calculus and other sample types, ordered by decreasing log-fold change. Bars represent standard error.

### 3.2 Samples show an increased mineralisation over the course of the experiment

Comparing the development of the model calculus to the reference samples, it is evident that between days 7 and 24 there is a decrease of the protein components and increase of the inorganic mineral carbonate hydroxyapatite. The model calculus samples from the end of the experiment are similar to both the modern and archaeological reference samples. The main difference is a lower organic component in reference samples seen as a reduced amide I peak at around 1637 compared to the carbonate peak at around 1420, and an absence of amide II and III. Further, there is a reduction in CH3 bands at 3000-2900 cm^−1^ (Figure 9A-D).

**Figure 9:**
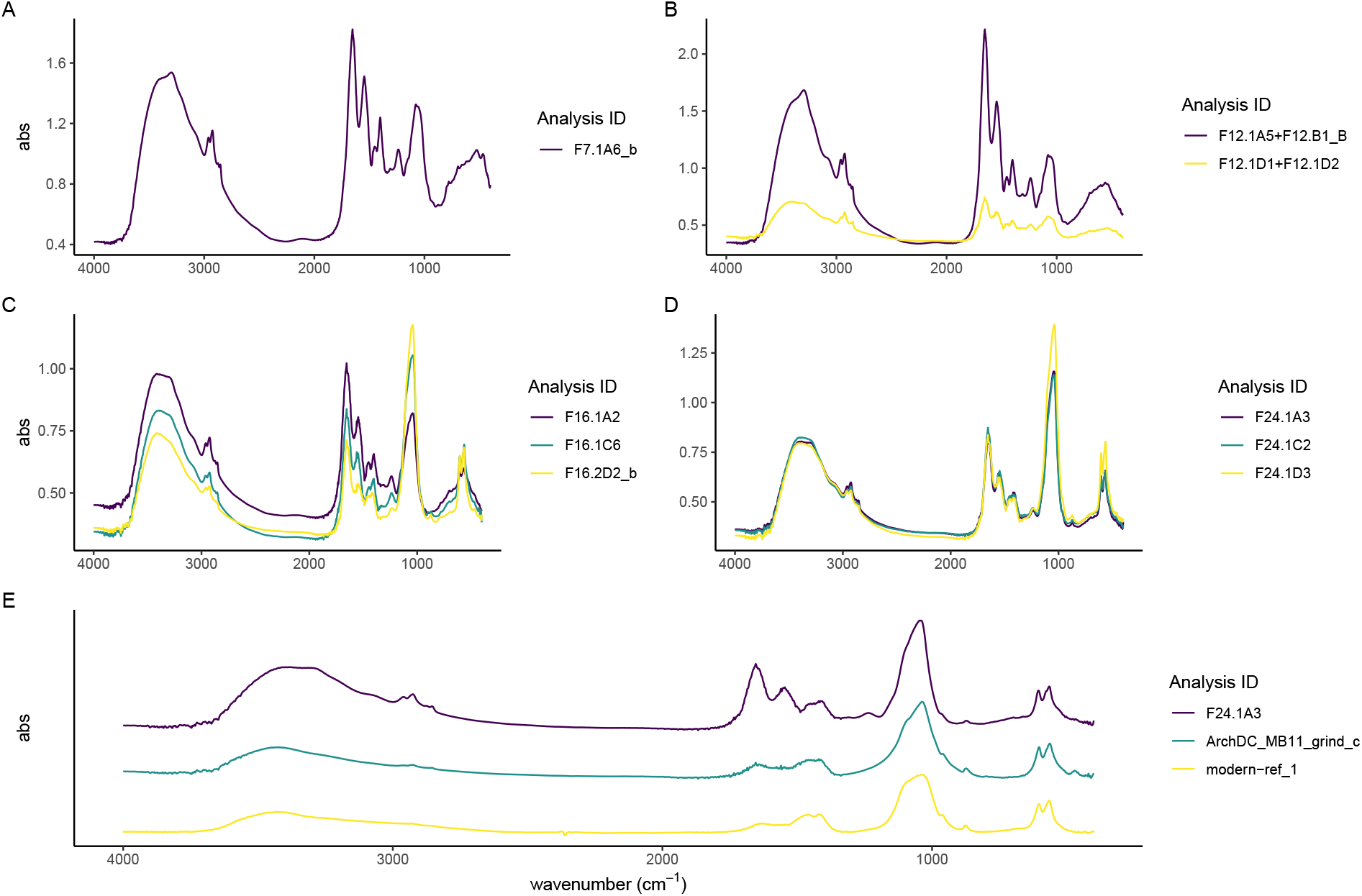
Select spectra from all experiment sampling days; (A) day 7, (B) day 12, (C) day 16, and (D) day 24. Absorbance bands in stretching mode around 3400 cm1 typical of the hydroxyl group. Analysis ID for model samples is constructed as: F[day sampled].[well sampled]_[grind sample].

Sample spectra from days 7 and 12 are characterised by a high content of proteins as evident by the strong amide I absorbance band at 1650, a less pronounced amide II band at 1545 cm^−1^, and the small amide III band at 1237 cm^−1^. Related to the organic component of the samples are also the three marked CH_3_ and CH_2_ stretching vibrations at 2960, 2920, and 2850 cm^−1^ wavenumbers. The presence of mineral component is evident from the presence of 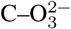 absorbance bands at 1450 and 1400 cm^−1^ wavenumbers typical of carbonates, and 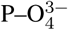 absorbance band at 1080 and 1056 cm^−1^ which are related to phosphate minerals. There is a large variation between the spectra, possibly indicating different formation rates of the different components in the samples (Figure 9A and B).

In spectra from days 16 to 24, the ratio of amides to PO_4_ has shifted, with the main peak shifting to the PO_4_ v_3_ absorbance band at 1039–1040 cm^−1^, indicating that the main component of the samples is carbonate hydroxyapatite. A well-defined PO_4_ doublet at 600 and 560 is present. Small 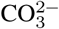 asymmetric stretching at 1450 cm^−1^ and 1415 cm^−1^, and stretching vibrations at 875-870 cm^−1^ indicate that the carbonate minerals component is also becoming more crystallised. There is a decreased variability between the spectra, with most spectra exhibiting a higher phosphate-to-protein/lipid ratio (Figure 9C and D).

### 3.3 Model calculus has a similar mineral composition to natural calculus

To determine whether the model dental calculus is comparable to natural dental calculus, both modern and archaeological dental calculus were analysed with FTIR spectroscopy to ascertain their composition. The archaeological and modern reference spectra are largely indistinguishable and consist of a broad O–H absorbance band (3400 cm^−1^) related to amid a and hydroxyl group, weak CH3 bands (3000–2900 cm^−1^), amide I band (1650 cm^−1^) which is related to the protein content, carbonate (1420, 1458-1450, 875-870 cm^−1^), and phosphates (1036-1040, 602-4, 563-566 cm^−1^) (Figure 9E) which, together with the hydroxyl and the carbonate, can be identified as derived from carbonate hydroxyapatite, the main mineral found in mature dental calculus (Hayashizaki et al., 2008; Jin & Yip, 2002).

### 3.4 Samples show similar crystallinity and order to reference calculus

We determined the level of crystallinity and order of the carbonate hydroxyapatite in our samples as an indication for its maturity by using the grinding curves method presented by Asscher, Regev, et al. (2011) and Asscher, Weiner, et al. (2011).

Samples were compared to published trendlines for archaeological and modern enamel (Asscher, Regev, et al., 2011). We see no appreciable differences between days 16, 20, and 24. The archaeological dental calculus shows a slightly increased slope compared to model calculus from the three sampling days used in the grind curve (Figure 10), possibly indicating larger crystal size due to more complete crystalisation. The steeper slope of enamel samples is consistent with a more ordered structure in enamel compared to dental calculus.

**Figure 10:**
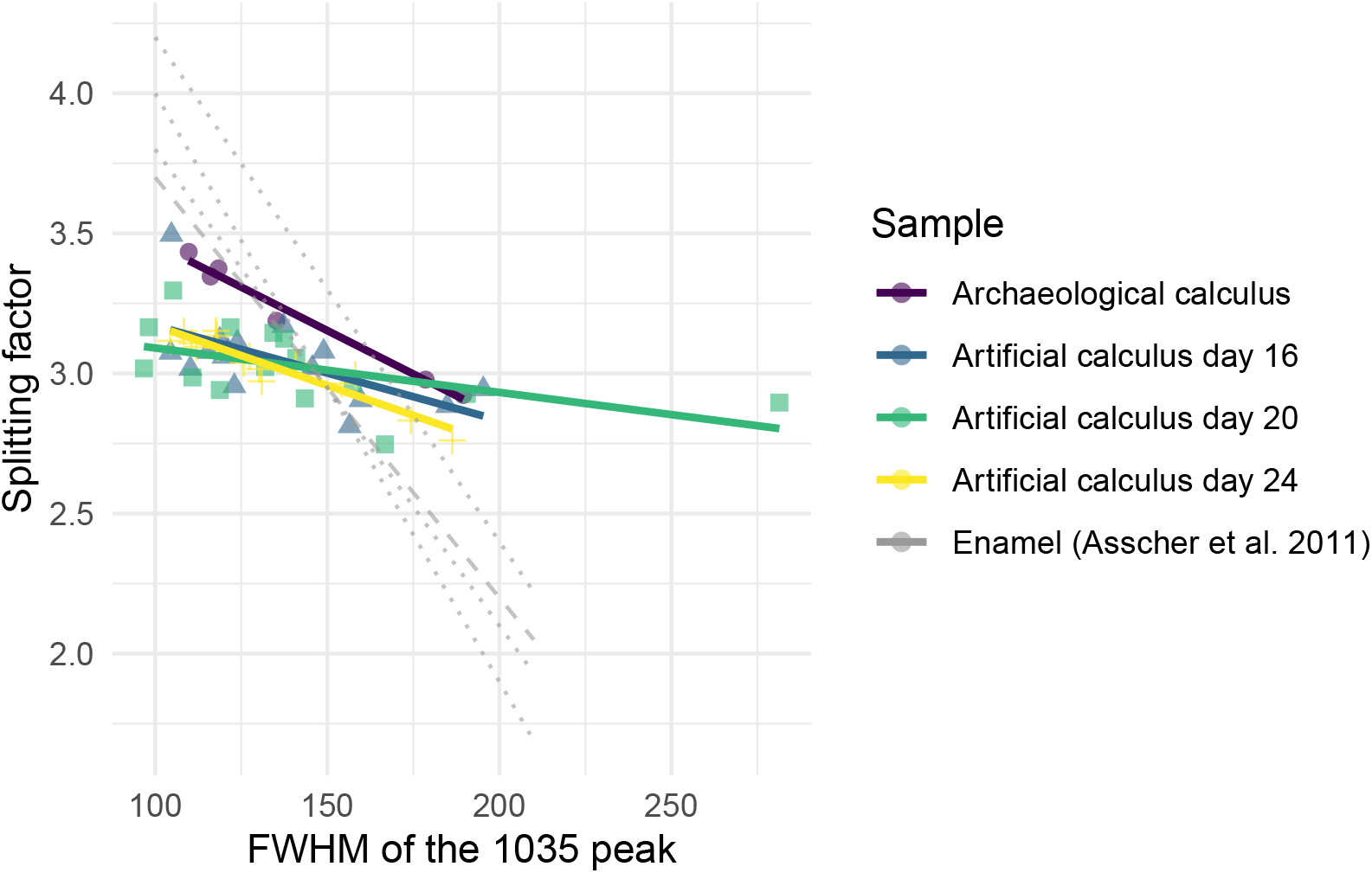
Grinding curves of our biofilm and model calculus compared to published trendlines (dashed light grey lines) for archaeological (dotted line) and modern (dashed line) enamel.

## 4 Discussion

In this study we present a calcifying oral biofilm model to produce artificial dental calculus. Our proposed use of the model is to address a variety of research questions related to dietary information extracted from dental calculus, in both modern and archaeological samples. For that to be feasible, the model needs to serve as a viable proxy to dental calculus grown under natural conditions, i.e., in the human oral cavity. It needs, as much as possible, to mimic the diversity and complexity of the natural oral microbiome, while also offering control over factors such as dietary input, growth conditions, and replicability within and between experiments. Here, we assessed the viability of our model as a proxy for dental calculus using metagenomic classification and FTIR analysis to explore the bacterial and mineral composition, and compare with oral reference samples.

### 4.1 Microbiome

Model calculus has lower species diversity than inocula saliva and oral reference samples, which is a common limitation in biofilm models (Bjarnsholt et al., 2013; Edlund et al., 2013). The donated saliva for the experiment had a lower diversity than the reference saliva samples, and may have contributed to a lower diversity in experimental samples. Consequently, there is also a lower diversity and richness when compared to other modern oral reference samples, including oral mucosa, saliva, plaque, and calculus. Samples of the medium from early in the experiment have similar species profiles to the donated saliva, but gradually diverge over the course of the experiment. This may be caused by experimental setup not sufficiently mimicking the oral environment, allowing species to thrive that do not normally thrive in the natural oral environment.

Oral reference samples have a relative abundance of streptococci similar to our model, but a more diverse representation from other genera and an overall higher species diversity and richness than our model. Reference samples also had a more diverse aerotolerance profile than our model, which primarily consisted of (faculatative) anaerobes. Species within predominantly aerobic genera, are deficient in the model, suggesting a shift from a largely aerotolerant profile to an anaerobic profile during the experiment. While our model is not set up as an anaerobic system, the anaerobes seem to have outcompeted aerobes and, to some extent, facultative anaerobes. This is likely a result of communities of bacteria within the biofilm creating favourable microenvironments facilitated by the protective properties of the biofilm matrix (Edlund et al., 2018; Flemming et al., 2016).

Overall, the majority of model calculus samples contained a distinctly oral signature, providing a promising starting point for the use of the model as a viable proxy to dental calculus. The main differences between model and oral reference samples may be due to human variation, as there can be large differences in the oral microbiome of two individuals at the species level due to variations in age, sex, and other demographic factors, as well as how and when saliva samples were collected (Burcham et al., 2020; Nearing et al., 2020). Whether or not distinct microbial profiles, and the extracellular matrix they produce, will affect the retention of dietary particles in plaque remains to be seen, but is an important question to address in the future.

### 4.2 Mineralisation

FTIR analysis allowed us to address the mineralisation process of the model, which showed an increasing mineral composition over the course of the experiment. As the model biofilm matured, the predominantly organic content of early samples was replaced by inorganic content in the form of carbonated hydroxyapatite, consistent with a shift from a high presence of bacterial cells in a matrix of extracellular polysaccharides (Jain et al., 2013; Sutherland, 2001; Zhang et al., 1998) to a predominantly mineral content.

The model calculus samples resemble both the modern reference calculus and the archaeological calculus in mineral composition and crystallinity. The steeper slope in the grind curve plots of the archaeological sample suggests that the crystals in archaeological samples are larger, and hence more ordered than in model calculus. A possible explanation is that the inorganic crystals within archaeological calculus have had more time to grow into the space left by degraded organic matter (Weiner, 2010a); however, we only analysed one archaeological sample and cannot definitively address this. The short duration of model calculus growth may also have affected the results, compared to the longer-term growth and mineralisation of natural calculus. The constant disruptions in growth of *in vivo* dental plaque/calculus, due to oral hygiene and other external pressures on biofilm growth, may lead to multiple stages of calcium phosphates, whereas our model has more stable growth conditions.

One of the most well-known biomineralisers, *Corynebacterium matruchotii* (Ennever et al., 1978; Takazoe et al., 1970), exhibited a lower abundance in our model calculus compared to modern reference calculus. However, the mineral composition of the end results were similar, reinforcing the idea that, under the right circumstances, biofilms with a range of microbial profiles can facilitate mineralisation (Moorer et al., 1993). Bacteria and their ability to secrete an extracellular matrix are integral in the formation of dental calculus, and inevitably serve as part of the structure that dental calculus is built upon (Rohanizadeh & LeGeros, 2005), while the exact species composition of the biofilm communities may be less important.

### 4.3 Replicability

Model calculus displayed similar species diversity and microbial profiles across all samples, indicating a high level of replicability between samples in the experimental run. It remains to be seen whether the replicability within the experiment also scales up to between-experiment replicability in our model, though others have already shown that replicability in long-term models is possible when using the same inocula (Velsko & Shaddox, 2018). The variation in mineral composition in our model was initially high, but samples from day 24 were largely similar in composition as observed in the FTIR spectra. The use of a simple multiwell plate setup allows us to submit many samples to the same conditions, increasing replicability between samples (Exterkate et al., 2010).

### 4.4 Limitations

While our in vitro model calculus system provides reproducible and consistent artificial dental calculus for archaeological research, as demonstrated by the species composition and the mineralisation properties, we recognise the model has several limitations. Our single-donor approach may have affected the diversity of the model. The donated saliva from our study had a lower mean Shannon Index than other saliva samples. The lower diversity may be caused by only using one donor instead of pooling saliva from multiple individuals. However, having a single inoculum donor allows us to maintain the integrity of a native oral microbiome which may be lost when samples are pooled (Edlund et al., 2013). It is also possible that the diversity was affected by the collection and storage methods we used. This has been shown to have minimal effect on microbial profiles at the genus level (Lim et al., 2017), but some effect on beta diversity calculations (Omori et al., 2021).

Some samples were grown with starch-sucrose solutions as nutrients, while controls were grown with sucrose only. Due to the financial cost, we did not sequence enough samples of each nutrient treatment to assess the influence of starch on the microbial community or mineral composition. Biofilms were grown in a standard shaking bacterial growth incubator, rather than an incubator specific to cell cultures. The lack of complex environmental control may cause the model to deviate from its natural growth over the 25 days that the experiment is run, due to a lack of precise control over conditions such as pH and salivary flow rates.

There is also the possibility that contamination was introduced into the model during the experiment. While the CPMU solution was prepared under sterile conditions, the solution itself was not autoclaved or filter-sterilised. In the species composition metagenomic analysis, all medium samples collected after the introduction of CPMU on day 14 were removed by the authentication step because the majority of species appeared to derive from environmental sources indicating external contamination. Going forward we recommend filter-sterilising solutions that are not autoclaved.

To avoid disturbing the growth and development of our biofilm, we took samples of media from the bottom of the wells after three days without full media replacement, careful not to disturb other plate-bound biofilms. The samples may therefore not fully reflect the composition of the biofilm itself. Going forward we recommend sampling from the actual biofilm, as this is the sample type under investigation.

### 4.5 Future work

Further protocol optimisation will also be necessary to address some of the limitations of our current model, such as reducing the frequency of medium replacement (currently every three days) to help promote the growth of slow-growing fastidious organisms and limit generalists such as enterococci, and supplementing it with serum to provide additional nutrients and biofilm stability (Ammann et al., 2012; Tian et al., 2010). More infrequent medium replacement would facilitate slow-growing bacteria in establishing their metabolic relationships, allowing the byproducts of some species to become abundant enough for others that depend on these to grow (Marsh, 2005).

Our goals for additional validation measures involve functional profiles of bacteria, to see if metabolic behaviour of bacteria is consistent with *in vivo* conditions, and whether this is affected by the presence/absence of amylase and starch treatments. The absence of host salivary *α*-amylase activity in our model (as shown in Bartholdy & Henry (2022)) provides an opportunity to explore the effect of various amylase levels on biofilm growth and composition, as well as the incorporation of dietary compounds in dental calculus.

The model can also be used to explore limitations and biases of methods used to reconstruct past dietary patterns from dental calculus. To this end, sucrose and raw starch treatments can be replaced with other dietary components of interest, such as cooked starches, whole plant extracts, and various proteins.

## 5 Conclusions

The bacterial profile of our model calculus is not an exact match to the natural modern or archaeological reference calculus, but species richness and diversity falls within a similar range as the reference *in vitro* model, and the core genera are predominantly oral. Our model calculus had a distinct microbial profile from modern reference calculus, but a similar mineral composition to modern and archaeological reference calculus, consisting of carbonate hydroxyapatite and similar levels of crystallinity and order, with a slightly higher organic phase.

Our model has many potential benefits within archaeological research, especially since the setup does not require highly specialised equipment, making it accessible to many labs within the archaeological sciences. It can be used to test many fundamental aspects of the process of incorporation, retention, and subsequent extraction of various dietary components from archaeological dental calculus. Using an oral biofilm model in a controlled environment with known dietary input, we can learn more about how different methods of food processing in the past may affect results of dental calculus analyses, and how the methods we use may further distort this picture. Our method can be used to test methods (e.g. DNA, proteomics, etc.), decontamination protocols, as well as training on these methods and protocols without depleting limited archaeological resources. The purpose of our model is not to replace studies conducted on archaeological and natural dental calculus, but rather to balance limitations of each method and serve as a complementary approach to expand our toolkit.

## Acknowledgements

We would like to thank the nf-core/eager community for assistance with the EAGER pipeline, especially Dr. James Fellows Yates. We also thank Sophie Seng for involvement in the DNA extraction. This work was performed using the compute resources from the Academic Leiden Interdisciplinary Cluster Environment (ALICE) provided by Leiden University, with special thanks to Dr. Robert Schulz. The FTIR analysis was conducted at the Laboratory for Sedimentary Archaeology, Haifa University, courtesy of Prof. Ruth Shahack-Gross, with additional help from Dr. Yotam Asscher.

This research has received funding from the European Research Council under the European Union’s Horizon 2020 research and innovation program, grant agreement number STG–677576 (“HARVEST”, funding BPB and AGH), as well as STG-948365 (“PROSPER”, funding ZF), and Werner Siemens Foundation (“PALEOBIOTECHNOLOGY, funding IMV and CW).

## Data Availability Statement

Human-filtered DNA sequencing data have been deposited in the ENA database under accession PRJEB61886. Analysis scripts and source code for the manuscript and supplementary materials are available on GitHub (https://github.com/bbartholdy/byoc-valid) and archived on 4TU.ResearchData (10.4121/99932661-fe79-4f4e-a812-a8917ad18fd0). FTIR data and spectra are available on 4TU.ResearchData (10.4121/466b2588-9689-4d84-a8a0-5216aa39e40b). Detailed protocols for growing the oral biofilm are available on protocols.io (10.17504/protocols.io.dm6gpj9rdgzp/v1).

## Notes

### Competing Interest Statement

The authors have declared no competing interest.

### Summary of Updates

Addition of author and funding information. Correct version of supplementary material uploaded.

## References

Adler, C. J., Dobney, K., Weyrich, L. S., Kaidonis, J., Walker, A. W., Haak, W., Bradshaw, C. J., Townsend, G., Sotysiak, A., Alt, K. W., Parkhill, J., & Cooper, A. (2013). Sequencing ancient calcified dental plaque shows changes in oral microbiota with dietary shifts of the Neolithic and Industrial revolutions. Nature Genetics, 45(4), 450–455, 455e1. https://doi.org/10.1038/ng.2536

Ammann, T. W., Gmür, R., & Thurnheer, T. (2012). Advancement of the 10-species subgingival Zurich biofilm model by examining different nutritional conditions and defining the structure of the in vitro biofilms. BMC Microbiology, 12, 227. https://doi.org/10.1186/1471-2180-12-227

Aron, F., Neumann, G., & Brandt, G. (2020). Half-UDG treated double-stranded ancient DNA library preparation for illumina sequencing v1 [Data set]. Protocols. Io.

Asscher, Y., Regev, L., Weiner, S., & Boaretto, E. (2011). Atomic Disorder in Fossil Tooth and Bone Mineral: An FTIR Study Using the Grinding Curve Method. ArcheoSciences. Revue d’archéométrie, 35, 35, 135–141. https://doi.org/10.4000/archeosciences.3062

Asscher, Y., Weiner, S., & Boaretto, E. (2011). Variations in Atomic Disorder in Biogenic Carbonate Hydroxyapatite Using the Infrared Spectrum Grinding Curve Method. Advanced Functional Materials, 21(17), 3308–3313. https://doi.org/10.1002/adfm.201100266

Bartholdy, B. P., Hasselstrøm, J. B., Sørensen, L. K., Casna, M., Hoogland, M., Beemster, H. G., & Henry, A. G. (2023). Multiproxy analysis exploring patterns of diet and disease in dental calculus and skeletal remains from a 19th century Dutch population. Zenodo. https://doi.org/10.5281/zenodo.7649151

Bartholdy, B. P., & Henry, A. G. (2022). Investigating Biases Associated With Dietary Starch Incorporation and Retention With an Oral Biofilm Model. Frontiers in Earth Science, 10. https://www.frontiersin.org/articles/10.3389/feart.2022.886512

Bjarnsholt, T., Alhede, M., Alhede, M., Eickhardt-Sørensen, S. R., Moser, C., Kühl, M., Jensen, P. Ø., & Høiby, N. (2013). The in vivo biofilm. Trends in Microbiology, 21(9), 466–474. https://doi.org/10.1016/j.tim.2013.06.002

Buckley, S., Usai, D., Jakob, T., Radini, A., & Hardy, K. (2014). Dental Calculus Reveals Unique Insights into Food Items, Cooking and Plant Processing in Prehistoric Central Sudan. PLOS ONE, 9(7), e100808. https://doi.org/10.1371/journal.pone.0100808

Burcham, Z. M., Garneau, N. L., Comstock, S. S., Tucker, R. M., Knight, R., Metcalf, J. L., Genetics of Taste Lab Citizen Scientists, Miranda, A., Reinhart, B., Meyers, D., Woltkamp, D., Boxer, E., Hutchens, J., Kim, K., Archer, M., McAteer, M., Huss, P., Defonseka, R., Stahle, S., … Reusser, W. (2020). Patterns of Oral Microbiota Diversity in Adults and Children: A Crowdsourced Population Study. Scientific Reports, 10(1), 2133. https://doi.org/10.1038/s41598-020-59016-0

Davis, N. M., Proctor, D. M., Holmes, S. P., Relman, D. A., & Callahan, B. J. (2018). Simple statistical identification and removal of contaminant sequences in marker-gene and metagenomics data. Microbiome, 6(1), 226. https://doi.org/10.1186/s40168-018-0605-2

Edlund, A., Yang, Y., Hall, A. P., Guo, L., Lux, R., He, X., Nelson, K. E., Nealson, K. H., Yooseph, S., Shi, W., & McLean, J. S. (2013). An in vitrobiofilm model system maintaining a highly reproducible species and metabolic diversity approaching that of the human oral microbiome. Microbiome, 1(1), 25. https://doi.org/10.1186/2049-2618-1-25

Edlund, A., Yang, Y., Yooseph, S., He, X., Shi, W., & McLean, J. S. (2018). Uncovering complex microbiome activities via metatranscriptomics during 24 hours of oral biofilm assembly and maturation. Microbiome, 6(1), 217. https://doi.org/10.1186/s40168-018-0591-4

Eerkens, J. W., Tushingham, S., Brownstein, K. J., Garibay, R., Perez, K., Murga, E., Kaijankoski, P., Rosenthal, J. S., & Gang, D. R. (2018). Dental calculus as a source of ancient alkaloids: Detection of nicotine by LC-MS in calculus samples from the Americas. Journal of Archaeological Science: Reports, 18, 509–515. https://doi.org/10.1016/j.jasrep.2018.02.004

Ennever, J., Riggan, L. J., Vogel, J. J., & Boyan-Salyers, B. (1978). Characterization of Bacterionema matruchotii Calcification Nucleator. Journal of Dental Research, 57(4), 637–642. https://doi.org/10.1177/00220345780570041901

Exterkate, R. A. M., Crielaard, W., & Ten Cate, J. M. (2010). Different Response to Amine Fluoride by Streptococcus mutans and Polymicrobial Biofilms in a Novel High-Throughput Active Attachment Model. Caries Research, 44(4), 372–379. https://doi.org/10.1159/000316541

Fagernäs, Z., Salazar-García, D. C., Haber Uriarte, M., Avilés Fernández, A., Henry, A. G., Lomba Maurandi, J., Ozga, A. T., Velsko, I. M., & Warinner, C. (2022). Understanding the microbial biogeography of ancient human dentitions to guide study design and interpretation. FEMS Microbes, 3, xtac006. https://doi.org/10.1093/femsmc/xtac006

Fellows Yates, J. A., Lamnidis, T. C., Borry, M., Valtueña, A. A., Fagernäs, Z., Clayton, S., Garcia, M. U., Neukamm, J., & Peltzer, A. (2020). Reproducible, portable, and efficient ancient genome reconstruction with nf-core/eager. bioRxiv, 2020.06.11.145615. https://doi.org/10.1101/2020.06.11.145615

Fellows Yates, J. A., Velsko, I. M., Aron, F., Posth, C., Hofman, C. A., Austin, R. M., Parker, C. E., Mann, A. E., Nägele, K., Arthur, K. W., Arthur, J. W., Bauer, C. C., Crevecoeur, I., Cupillard, C., Curtis, M. C., Dalén, L., Bonilla, M. D.-Z., Fernández-Lomana, J. C. D., Drucker, D. G., … Warinner, C. (2021). The evolution and changing ecology of the African hominid oral microbiome. Proceedings of the National Academy of Sciences, 118(20). https://doi.org/10.1073/pnas.2021655118

Filoche, S. K., Soma, K. J., & Sissons, C. H. (2007). Caries-related plaque microcosm biofilms developed in microplates. Oral Microbiology and Immunology, 22(2), 73–79. https://doi.org/10.1111/j.1399-302X.2007.00323.x

Flemming, H.-C., Wingender, J., Szewzyk, U., Steinberg, P., Rice, S. A., & Kjelleberg, S. (2016). Biofilms: An emergent form of bacterial life. Nature Reviews Microbiology, 14(9), 563–575. https://doi.org/10.1038/nrmicro.2016.94

Gloor, G. B., Macklaim, J. M., Pawlowsky-Glahn, V., & Egozcue, J. J. (2017). Microbiome Datasets Are Compositional: And This Is Not Optional. Frontiers in Microbiology, 8, 2224. https://doi.org/10.3389/fmicb.2017.02224

Hardy, K., Blakeney, T., Copeland, L., Kirkham, J., Wrangham, R., & Collins, M. (2009). Starch granules, dental calculus and new perspectives on ancient diet. Journal of Archaeological Science, 36(2), 248–255. https://doi.org/10.1016/j.jas.2008.09.015

Hayashizaki, J., Ban, S., Nakagaki, H., Okumura, A., Yoshii, S., & Robinson, C. (2008). Site specific mineral composition and microstructure of human supra-gingival dental calculus. Archives of Oral Biology, 53(2), 168–174. https://doi.org/10.1016/j.archoralbio.2007.09.003

Hendy, J., Warinner, C., Bouwman, A., Collins, M. J., Fiddyment, S., Fischer, R., Hagan, R., Hofman, C. A., Holst, M., Chaves, E., Klaus, L., Larson, G., Mackie, M., McGrath, K., Mundorff, A. Z., Radini, A., Rao, H., Trachsel, C., Velsko, I. M., & Speller, C. F. (2018). Proteomic evidence of dietary sources in ancient dental calculus. Proceedings. Biological Sciences, 285(1883), 20180977. https://doi.org/10.1098/rspb.2018.0977

Henry, A. G., & Piperno, D. R. (2008). Using plant microfossils from dental calculus to recover human diet: A case study from Tell al-Raq’i, Syria. Journal of Archaeological Science, 35(7), 1943–1950. https://doi.org/10.1016/j.jas.2007.12.005

Jain, K., Parida, S., Mangwani, N., Dash, H. R., & Das, S. (2013). Isolation and characterization of biofilm-forming bacteria and associated extracellular polymeric substances from oral cavity. Annals of Microbiology, 63(4), 1553– 1562. https://doi.org/10.1007/s13213-013-0618-9

Ji, H., Nakagaki, H., Hayashizaki, J., Tsuboi, S., Kato, K., Toyama, A., Arai, K., Thuy, T. T., Ha, N. T. T., Kameyama, Y., Kirkham, J., & Robinson, C. (2000). Fluoride and magnesium concentrations in human dental calculus obtained from Japanese and Chinese patients. Archives of Oral Biology, 45(7), 611–615. https://doi.org/10.1016/S0003-9969(00)00021-2

Jin, Y., & Yip, H.-K. (2002). Supragingival Calculus: Formation and Control. Critical Reviews in Oral Biology & Medicine. https://doi.org/10.1177/154411130201300506

Kazarina, A., Petersone-Gordina, E., Kimsis, J., Kuzmicka, J., Zayakin, P., Grikjans, ., Gerhards, G., & Ranka, R. (2021). The Postmedieval Latvian Oral Microbiome in the Context of Modern Dental Calculus and Modern Dental Plaque Microbial Profiles. Genes, 12(2), 309. https://doi.org/10.3390/genes12020309

Knights, D., Kuczynski, J., Charlson, E. S., Zaneveld, J., Mozer, M. C., Collman, R. G., Bushman, F. D., Knight, R., & Kelley, S. T. (2011). Bayesian community-wide culture-independent microbial source tracking. Nature Methods, 8(9), 761–763. https://doi.org/10.1038/nmeth.1650

Lemmers, S. A. M., Schats, R., Hoogland, M. L. P., & Waters-Rist, A. (2013). Fysisch antropologische analyse Middenbeemster. In De begravingen bij de Keyserkerk te Middenbeemster (pp. 35–60).

Leonard, C., Vashro, L., O’Connell, J. F., & Henry, A. G. (2015). Plant microremains in dental calculus as a record of plant consumption: A test with Twe forager-horticulturalists. Journal of Archaeological Science: Reports, 2, 449–457. https://doi.org/10.1016/j.jasrep.2015.03.009

Li, H., & Durbin, R. (2009). Fast and accurate short read alignment with Burrows–Wheeler transform. Bioinformatics, 25(14), 1754–1760. https://doi.org/10.1093/bioinformatics/btp324

Lim, Y., Totsika, M., Morrison, M., & Punyadeera, C. (2017). The saliva microbiome profiles are minimally affected by collection method or DNA extraction protocols. Scientific Reports, 7(1, 1), 8523. https://doi.org/10.1038/s41598-017-07885-3

Lin, H., & Peddada, S. D. (2020). Analysis of compositions of microbiomes with bias correction. Nature Communications, 11(1, 1), 3514. https://doi.org/10.1038/s41467-020-17041-7

Ma, Z., Liu, S., Li, Z., Ye, M., & Huan, X. (2022). Human Diet Patterns During the Qijia Cultural Period: Integrated Evidence of Stable Isotopes and Plant Micro-remains From the Lajia Site, Northwest China. Frontiers in Earth Science, 10. https://www.frontiersin.org/articles/10.3389/feart.2022.884856

Marsh, P. D. (2005). Dental plaque: Biological significance of a biofilm and community life-style. Journal of Clinical Periodontology, 32(6), 7–15. https://doi.org/10.1111/j.1600-051X.2005.00790.x

Marsh, P. D. (2006). Dental plaque as a biofilm and a microbial community – implications for health and disease. BMC Oral Health, 6(S1), S14. https://doi.org/10.1186/1472-6831-6-S1-S14

Mentzer, S. M., Miller, C. E., Kloos, P., Wadley, L., & Conard, N. J. (2014). The distribution of authigenic minerals in the Middle Stone Age deposits of Sibudu (South Africa), and implications for the preservation of archaeological features. European Society for the Study of Human Evolution, 4thAnnual Meeting, Florence, Italy.

Mickleburgh, H. L., & Pagán-Jiménez, J. R. (2012). New insights into the consumption of maize and other food plants in the pre-Columbian Caribbean from starch grains trapped in human dental calculus. Journal of Archaeological Science, 39(7), 2468–2478. https://doi.org/10.1016/j.jas.2012.02.020

Middleton, J. D. (1965). Human salivary proteins and artificial calculus formation in vitro. Archives of Oral Biology, 10(2), 227–235. https://doi.org/10.1016/0003-9969(65)90024-5

Moorer, W. R., Ten Cate, J. M., & Buijs, J. F. (1993). Calcification of a Cariogenic Streptococcus and of Corynebacterium (Bacterionema) matruchotii. Journal of Dental Research, 72(6), 1021–1026. https://doi.org/10.1177/00220345930720060501

Nearing, J. T., DeClercq, V., Van Limbergen, J., & Langille, M. G. I. (2020). Assessing the Variation within the Oral Microbiome of Healthy Adults. mSphere, 5(5), e00451–20. https://doi.org/10.1128/mSphere.00451-20

Oksanen, J., Simpson, G. L., Blanchet, F. G., Kindt, R., Legendre, P., Minchin, P. R., O’Hara, R. B., Solymos, P., Stevens, M. H. H., Szoecs, E., Wagner, H., Barbour, M., Bedward, M., Bolker, B., Borcard, D., Carvalho, G., Chirico, M., De Caceres, M., Durand, S., … Weedon, J. (2022). Vegan: Community ecology package [Manual]. https://CRAN.R-project.org/package=vegan

Omelon, S., Ariganello, M., Bonucci, E., Grynpas, M., & Nanci, A. (2013). A Review of Phosphate Mineral Nucleation in Biology and Geobiology. Calcified Tissue International, 93(4), 382–396. https://doi.org/10.1007/s00223-013-9784-9

Omori, M., Kato-Kogoe, N., Sakaguchi, S., Fukui, N., Yamamoto, K., Nakajima, Y., Inoue, K., Nakano, H., Motooka, D., Nakano, T., Nakamura, S., & Ueno, T. (2021). Comparative evaluation of microbial profiles of oral samples obtained at different collection time points and using different methods. Clinical Oral Investigations, 25(5), 2779– 2789. https://doi.org/10.1007/s00784-020-03592-y

Pearce, E. I. F., & Sissons, C. H. (1987). The Concomitant Deposition of Strontium and Fluoride in Dental Plaque. Journal of Dental Research, 66(10), 1518–1522. https://doi.org/10.1177/00220345870660100101

Power, R. C., Salazar-Garcia, D. C., Wittig, R. M., Freiberg, M., & Henry, A. G. (2015). Dental calculus evidence of Tai Forest Chimpanzee plant consumption and life history transitions. Scientific Reports, 5, 15161. https://doi.org/10.1038/srep15161

R Core Team. (2020). R: A language and environment for statistical computing [Manual]. R Foundation for Statistical Computing; R Foundation for Statistical Computing. https://www.R-project.org/

Radini, A., & Nikita, E. (2022). Beyond dirty teeth: Integrating dental calculus studies with osteoarchaeological parameters. Quaternary International. https://doi.org/10.1016/j.quaint.2022.03.003

Reimer, L. C., Sardà Carbasse, J., Koblitz, J., Ebeling, C., Podstawka, A., & Overmann, J. (2022). BacDive in 2022: The knowledge base for standardized bacterial and archaeal data. Nucleic Acids Research, 50(D1), D741–D746. https://doi.org/10.1093/nar/gkab961

Rohanizadeh, R., & LeGeros, R. Z. (2005). Ultrastructural study of calculus–enamel and calculus–root interfaces. Archives of Oral Biology, 50(1), 89–96. https://doi.org/10.1016/j.archoralbio.2004.07.001

Rohart, F., Gautier, B., Singh, A., & Le Cao, K.-A. (2017). mixOmics: An R package for ‘omics feature selection and multiple data integration. PLoS Computational Biology, 13(11), e1005752. http://www.mixOmics.org

Schubert, M., Lindgreen, S., & Orlando, L. (2016). AdapterRemoval v2: Rapid adapter trimming, identification, and read merging. BMC Research Notes, 9, 88. https://doi.org/10.1186/s13104-016-1900-2

Shellis, R. P. (1978). A synthetic saliva for cultural studies of dental plaque. Archives of Oral Biology, 23(6), 485–489. https://doi.org/10.1016/0003-9969(78)90081-X

Sissons, C. H., Cutress, T. W., Hoffman, M. P., & Wakefield, J. S. J. (1991). A Multi-station Dental Plaque Microcosm (Artificial Mouth) for the Study of Plaque Growth, Metabolism, pH, and Mineralization: Journal of Dental Research. https://doi.org/10.1177/00220345910700110301

Stahl, R., Warinner, C., Velsko, I., Orfanou, E., Aron, F., & Brandt, G. (2019). Illumina double-stranded DNA dual indexing for ancient DNA v1 [Data set]. Protocols. Io.

Sutherland, I. W. (2001). The biofilm matrix – an immobilized but dynamic microbial environment. Trends in Microbiology, 9(5), 222–227. https://doi.org/10.1016/S0966-842X(01)02012-1

Takazoe, I., Vogel, J., & Ennever, J. (1970). Calcium Hydroxyapatite Nucleation by Lipid Extract of Bacterionema matruchotii. Journal of Dental Research, 49(2), 395–398. https://doi.org/10.1177/00220345700490023301

Tian, Y., He, X., Torralba, M., Yooseph, S., Nelson, K. e., Lux, R., McLean, J. s., Yu, G., & Shi, W. (2010). Using DGGE profiling to develop a novel culture medium suitable for oral microbial communities. Molecular Oral Microbiology, 25(5), 357–367. https://doi.org/10.1111/j.2041-1014.2010.00585.x

Tønjum, T., & van Putten, J. (2017). 179 - Neisseria. In J. Cohen, W. G. Powderly, & S. M. Opal (Eds.), Infectious Diseases (Fourth Edition) (pp. 1553–1564.e1). Elsevier. https://doi.org/10.1016/B978-0-7020-6285-8.00179-9

Tromp, M., Buckley, H., Geber, J., & Matisoo-Smith, E. (2017). EDTA decalcification of dental calculus as an alternate means of microparticle extraction from archaeological samples. Journal of Archaeological Science: Reports, 14, 461–466. https://doi.org/10.1016/j.jasrep.2017.06.035

Velsko, I. M., Cruz-Almeida, Y., Huang, H., Wallet, S. M., & Shaddox, L. M. (2017). Cytokine response patterns to complex biofilms by mononuclear cells discriminate patient disease status and biofilm dysbiosis. Journal of Oral Microbiology, 9(1), 1330645. https://doi.org/10.1080/20002297.2017.1330645

Velsko, I. M., Fellows Yates, J. A., Aron, F., Hagan, R. W., Frantz, L. A. F., Loe, L., Martinez, J. B. R., Chaves, E., Gosden, C., Larson, G., & Warinner, C. (2019). Microbial differences between dental plaque and historic dental calculus are related to oral biofilm maturation stage. Microbiome, 7(1), 102. https://doi.org/10.1186/s40168-019-0717-3

Velsko, I. M., Overmyer, K. A., Speller, C., Klaus, L., Collins, M. J., Loe, L., Frantz, L. A. F., Sankaranarayanan, K., Lewis, C. M., Martinez, J. B. R., Chaves, E., Coon, J. J., Larson, G., & Warinner, C. (2017). The dental calculus metabolome in modern and historic samples. Metabolomics, 13(11), 134. https://doi.org/10.1007/s11306-017-1270-3

Velsko, I. M., & Shaddox, L. M. (2018). Consistent and reproducible long-term in vitro growth of health and disease-associated oral subgingival biofilms. BMC Microbiology, 18(1), 70. https://doi.org/10.1186/s12866-018-1212-x

Warinner, C., Rodrigues, J. F., Vyas, R., Trachsel, C., Shved, N., Grossmann, J., Radini, A., Hancock, Y., Tito, R. Y., Fiddyment, S., Speller, C., Hendy, J., Charlton, S., Luder, H. U., Salazar-Garcia, D. C., Eppler, E., Seiler, R., Hansen, L. H., Castruita, J. A., … Cappellini, E. (2014). Pathogens and host immunity in the ancient human oral cavity. Nature Genetics, 46(4), 336–344. https://doi.org/10.1038/ng.2906

Warinner, C., Speller, C., & Collins, M. J. (2015). A new era in palaeomicrobiology: Prospects for ancient dental calculus as a long-term record of the human oral microbiome. Philosophical Transactions of the Royal Society B: Biological Sciences, 370(1660), 20130376. https://doi.org/10.1098/rstb.2013.0376

Weiner, S. (2010a). Biological Materials: Bones and Teeth. In Microarchaeology: Beyond the Visible Archaeological Record (pp. 99–134). Cambridge University Press.

Weiner, S. (2010b). Infrared Spectroscopy in Archaeology. In Microarchaeology: Beyond the Visible Archaeological Record (1st ed., pp. 275–316). Cambridge University Press. https://doi.org/10.1017/CBO9780511811210

Weiner, S., & Bar-Yosef, O. (1990). States of preservation of bones from prehistoric sites in the Near East: A survey. Journal of Archaeological Science, 17(2), 187–196. https://doi.org/10.1016/0305-4403(90)90058-D

White, D. J. (1997). Dental calculus: Recent insights into occurrence, formation, prevention, removal and oral health effects of supragingival and subgingival deposits. European Journal of Oral Sciences, 105(5), 508–522. https://doi.org/10.1111/j.1600-0722.1997.tb00238.x

Wickham, H. (2016). Ggplot2: Elegant Graphics for Data Analysis. Springer-Verlag. https://ggplot2.tidyverse.org

Wickham, Hadley, Averick, M., Bryan, J., Chang, W., McGowan, L. D., François, R., Grolemund, G., Hayes, A., Henry, L., Hester, J., Kuhn, M., Pedersen, T. L., Miller, E., Bache, S. M., Müller, K., Ooms, J., Robinson, D., Seidel, D. P., Spinu, V., … Yutani, H. (2019). Welcome to the tidyverse. Journal of Open Source Software, 4(43), 1686. https://doi.org/10.21105/joss.01686

Wong, L., Sissons, C. H., Pearce, E. I. F., & Cutress, T. W. (2002). Calcium phosphate deposition in human dental plaque microcosm biofilms induced by a ureolytic pH-rise procedure. Archives of Oral Biology, 47(11), 779–790. https://doi.org/10.1016/S0003-9969(02)00114-0

Wood, D. E., Lu, J., & Langmead, B. (2019). Improved metagenomic analysis with Kraken 2. Genome Biology, 20(1), 257. https://doi.org/10.1186/s13059-019-1891-0

Zhang, X., Bishop, P. L., & Kupferle, M. J. (1998). Measurement of polysaccharides and proteins in biofilm extracellular polymers. Water Science and Technology, 37(4), 345–348. https://doi.org/10.1016/S0273-1223(98)00127-9

