## Supplementary Materials for "Assessing the validity of a calcifying oral biofilm model as a suitable proxy for dental calculus"

#### Samples

Samples taken for DNA sequencing and FTIR analysis. Samples for DNA were collected from a seperate experimental run than samples for DNA, but experimental conditions were the same in each. Samples for DNA were medium collected from the bottom of each well after three days of growth (before medium was refreshed). Samples for FTIR were taken directly from the biofilm and dried prior to analysis.

### DNA

Table 1: Table of biofilm samples from this study. Includes which day in the experiment the sample was taken, and sample type (Env).

| #SampleID | day | Env |
|---|---|---|
| SYN001.A0101 | 0 | saliva |
| SYN002.A0101 | 3 | saliva |
| SYN017.A0101 | 5 | medium |
| SYN014.A0101 | 5 | medium |
| SYN003.A0101 | 5 | saliva |
| SYN015.B0101 | 7 | medium |
| SYN017.B0101 | 7 | medium |
| SYN018.C0101 | 9 | medium |
| SYN014.C0101 | 9 | medium |
| SYN015.D0101 | 12 | medium |
| SYN017.D0101 | 12 | medium |
| SYN015.E0101 | 15 | medium |
| SYN017.E0101 | 15 | medium |
| SYN015.F0101 | 18 | medium |
| SYN017.F0101 | 18 | medium |
| SYN015.G0101 | 21 | medium |
| SYN017.G0101 | 21 | medium |
| SYN015.H0101 | 24 | medium |
| SYN018.H0101 | 24 | medium |
| SYN005.I0101 | 24 | model_calculus |
| SYN006.I0101 | 24 | model_calculus |
| SYN008.I0101 | 24 | model_calculus |
| SYN009.I0101 | 24 | model_calculus |
| SYN015.I0101 | 24 | model_calculus |
| SYN018.I0101 | 24 | model_calculus |
| SYN019.I0101 | 24 | model_calculus |
| SYN022.I0101 | 24 | model_calculus |
| SYN023.I0101 | 24 | model_calculus |
| SYN025.I0101 | 24 | model_calculus |
| SYN028.I0101 | 24 | model_calculus |
| SYN012.I0101 | 24 | model_calculus |
| SYN013.I0101 | 24 | model_calculus |
| SYN016.I0101 | 24 | model_calculus |
| SYN021.I0101 | 24 | model_calculus |
| SYN026.I0101 | 24 | model_calculus |

### FTIR

Table 2: Table of oral reference samples. Includes which day in the experiment the sample was taken, and sample type.

| sample_id | day | sample_type |
|---|---|---|
| F7.1A6 | 7 | model_biofilm |
| F7.2D1 | 7 | model_biofilm |
| F7.1C4 | 7 | model_biofilm |
| F12.1A5 | 12 | model_biofilm |
| F12.1D1 | 12 | model_biofilm |
| F12.1D2 | 12 | model_biofilm |
| F12.1B1 | 12 | model_biofilm |
| F16.1A2 | 16 | model_biofilm |
| F16.1B2 | 16 | model_biofilm |
| F16.1C6 | 16 | model_biofilm |
| F16.1D6 | 16 | model_biofilm |
| F16.2B2 | 16 | model_biofilm |
| F16.2C2 | 16 | model_biofilm |
| F16.2D2 | 16 | model_biofilm |
| F20.1A1 | 20 | model_biofilm |
| F20.1B5 | 20 | model_biofilm |
| F20.1D5 | 20 | model_biofilm |
| F20.2A5 | 20 | model_biofilm |
| F20.2C5 | 20 | model_biofilm |
| F20.2D5 | 20 | model_biofilm |
| F24.1A3 | 24 | model_calculus |
| F24.1B3 | 24 | model_calculus |
| F24.1C2 | 24 | model_calculus |
| F24.1D3 | 24 | model_calculus |
| F24.2A4 | 24 | model_calculus |
| F24.2B4 | 24 | model_calculus |
| F24.2C3 | 24 | model_calculus |
| F24.2D3 | 24 | model_calculus |
| ArchDC_MB11 | NA | calculus |
| modern-ref_1 | NA | calculus |
| modern-ref_2 | NA | calculus |

#### Reference database and sequences

The reference database used in the EAGER pipeline was the Standard Kraken 2 database, downloaded from https://genome-idx.s3.amazonaws.com/kraken/k2_standard_20210517.tar.gz

```
wget https://genome-idx.s3.amazonaws.com/kraken/k2_standard_20210517.tar.gz
```

The human reference genome GRCh38 was downloaded on 2022-04-22.

```
wget ftp://ftp.ncbi.nlm.nih.gov/refseq/H_sapiens/annotation/GRCh38_latest/refseq_identifie
```

Table 3: Table of oral reference samples. Includes sample type (Env), and associated project and study with DOI.

| #SampleID | Env | Project | study_doi |
|---|---|---|---|
| SRS052668 | buccal_mucosa | HMP | NA |
| SRS011247 | buccal_mucosa | HMP | NA |
| SRS056892 | buccal_mucosa | HMP | NA |
| SRS011310 | buccal_mucosa | HMP | NA |
| SRS012281 | buccal_mucosa | HMP | NA |
| SRS013711 | buccal_mucosa | HMP | NA |
| SRS023591 | buccal_mucosa | HMP | NA |
| SRS058105 | buccal_mucosa | HMP | NA |
| SRS063478 | buccal_mucosa | HMP | NA |
| SRS075406 | buccal_mucosa | HMP | NA |
| JAE006.A0101 | modern_calculus | PRJEB31185 | 10.1186/s40168-019-0717-3 |
| JAE007.A0101 | modern_calculus | PRJEB31185 | 10.1186/s40168-019-0717-3 |
| JAE008.A0101 | modern_calculus | PRJEB31185 | 10.1186/s40168-019-0717-3 |
| JAE009.A0101 | modern_calculus | PRJEB31185 | 10.1186/s40168-019-0717-3 |
| JAE010.A0101 | modern_calculus | PRJEB31185 | 10.1186/s40168-019-0717-3 |
| JAE012.A0101 | modern_calculus | PRJEB31185 | 10.1186/s40168-019-0717-3 |
| JAE013.A0101 | modern_calculus | PRJEB31185 | 10.1186/s40168-019-0717-3 |
| JAE014.A0101 | modern_calculus | PRJEB31185 | 10.1186/s40168-019-0717-3 |
| JAE015.A0101 | modern_calculus | PRJEB31185 | 10.1186/s40168-019-0717-3 |
| JAE016.A0101 | modern_calculus | PRJEB31185 | 10.1186/s40168-019-0717-3 |
| SRS019120 | saliva | HMP | NA |
| SRS014468 | saliva | HMP | NA |
| SRS015055 | saliva | HMP | NA |
| SRS014692 | saliva | HMP | NA |
| SRS013942 | saliva | HMP | NA |
| SRS014477 | subgingival_plaque | HMP | NA |

| #SampleID | Env | Project | study_doi |
|---|---|---|---|
| SRS063215 | subgingival_plaque | HMP | NA |
| SRS019129 | subgingival_plaque | HMP | NA |
| SRS013950 | subgingival_plaque | HMP | NA |
| SRS015064 | subgingival_plaque | HMP | NA |
| SRS019029 | subgingival_plaque | HMP | NA |
| SRS014107 | subgingival_plaque | HMP | NA |
| SRS014691 | subgingival_plaque | HMP | NA |
| SRS021960 | supragingival_plaque | HMP | NA |
| SRS045313 | supragingival_plaque | HMP | NA |
| SRS047265 | supragingival_plaque | HMP | NA |
| SRS051378 | supragingival_plaque | HMP | NA |
| SRS015755 | supragingival_plaque | HMP | NA |
| SRS015378 | supragingival_plaque | HMP | NA |
| SRS022149 | supragingival_plaque | HMP | NA |
| SRS064493 | supragingival_plaque | HMP | NA |
| SRS052604 | supragingival_plaque | HMP | NA |
| SRS019333 | supragingival_plaque | HMP | NA |
| SRR5975923 | vitro_biofilm | PRJNA398963 | 10.1186/s40168-018-0591-4 |
| SRR5975924 | vitro_biofilm | PRJNA398963 | 10.1186/s40168-018-0591-4 |
| SRR5975925 | vitro_biofilm | PRJNA398963 | 10.1186/s40168-018-0591-4 |
| SRR5975926 | vitro_biofilm | PRJNA398963 | 10.1186/s40168-018-0591-4 |
| SRR5975927 | vitro_biofilm | PRJNA398963 | 10.1186/s40168-018-0591-4 |
| SRR5975928 | vitro_biofilm | PRJNA398963 | 10.1186/s40168-018-0591-4 |
| SRR5975929 | vitro_biofilm | PRJNA398963 | 10.1186/s40168-018-0591-4 |
| SRR5975930 | vitro_biofilm | PRJNA398963 | 10.1186/s40168-018-0591-4 |
| SRR5975931 | vitro_biofilm | PRJNA398963 | 10.1186/s40168-018-0591-4 |
| SRR5975932 | vitro_biofilm | PRJNA398963 | 10.1186/s40168-018-0591-4 |
| SRR5975933 | vitro_biofilm | PRJNA398963 | 10.1186/s40168-018-0591-4 |
| SRR5975934 | vitro_biofilm | PRJNA398963 | 10.1186/s40168-018-0591-4 |
| SRR5975936 | vitro_biofilm | PRJNA398963 | 10.1186/s40168-018-0591-4 |
| SRR5975937 | vitro_biofilm | PRJNA398963 | 10.1186/s40168-018-0591-4 |

Table 4: Table of environmental reference samples (contamination testing). Includes sample type (Env), and associated project and study with DOI.

| #SampleID | Env | Project | study_doi |
|---|---|---|---|
| SRR5420280 | indoor_air | PRJNA422794 | NA |
| SRR5420282 | indoor_air | PRJNA422794 | NA |
| SRR6129806 | indoor_air | PRJNA422794 | NA |
| SRR6129807 | indoor_air | PRJNA422794 | NA |

| #SampleID | Env | Project | study_doi |
|---|---|---|---|
| SRR6129808 | indoor_air | PRJNA422794 | NA |
| SRR6129809 | indoor_air | PRJNA422794 | NA |
| SRR6129810 | indoor_air | PRJNA422794 | NA |
| SRR6129811 | indoor_air | PRJNA422794 | NA |
| ERR1883419 | sediment | PRJEB18629 | 10.1126/science.aam9695 |
| ERR1883420 | sediment | PRJEB18629 | 10.1126/science.aam9695 |
| ERR1883421 | sediment | PRJEB18629 | 10.1126/science.aam9695 |
| ERR1883422 | sediment | PRJEB18629 | 10.1126/science.aam9695 |
| ERR1883423 | sediment | PRJEB18629 | 10.1126/science.aam9695 |
| ERR1883424 | sediment | PRJEB18629 | 10.1126/science.aam9695 |
| ERR1883430 | sediment | PRJEB18629 | 10.1126/science.aam9695 |
| ERR1883436 | sediment | PRJEB18629 | 10.1126/science.aam9695 |
| ERR1883438 | sediment | PRJEB18629 | 10.1126/science.aam9695 |
| SRR1631060 | skin | PRJNA46333 | NA |
| SRR1631061 | skin | PRJNA46333 | NA |
| SRR1631063 | skin | PRJNA46333 | NA |
| SRR1631064 | skin | PRJNA46333 | NA |
| SRR1633008 | skin | PRJNA46333 | NA |
| SRR3184100 | skin | PRJNA46333 | NA |
| SRR3184876 | skin | PRJNA46333 | NA |
| SRR3189411 | skin | PRJNA46333 | NA |
| SRR3189416 | skin | PRJNA46333 | NA |
| SRR3189418 | skin | PRJNA46333 | NA |
| SRR059389 | stool | PRJNA48479 | NA |
| SRR059425 | stool | PRJNA48479 | NA |
| SRR059455 | stool | PRJNA48479 | NA |
| SRR059917 | stool | PRJNA48479 | NA |
| SRR060358 | stool | PRJNA48479 | NA |
| SRR1761677 | stool | PRJNA268964 | NA |
| SRR1761682 | stool | PRJNA268964 | NA |
| SRR1761688 | stool | PRJNA268964 | NA |
| SRR1761692 | stool | PRJNA268964 | NA |
| SRR1761697 | stool | PRJNA268964 | NA |
| SRR1761698 | stool | PRJNA268964 | NA |
| SRR1761705 | stool | PRJNA268964 | NA |
| SRR1761710 | stool | PRJNA268964 | NA |
| SRR1761718 | stool | PRJNA268964 | NA |
| SRR1761721 | stool | PRJNA268964 | NA |
| SRR1929408 | stool | PRJNA278393 | NA |
| SRR1930121 | stool | PRJNA278393 | NA |
| SRR1930123 | stool | PRJNA278393 | NA |

| #SampleID | Env | Project | study_doi |
|---|---|---|---|
| SRR1930141 | stool | PRJNA278393 | NA |
| SRR1930145 | stool | PRJNA278393 | NA |

### Pre-processing

#### SourceTracker2

Steps taken for SourceTracker analysis.

OTU table was filtered for relative abundance. Percent abundance of each taxon across all samples was calculated and then taxa with lower than 0.001% abundance were filtered out.

Installation of Qiime2 and dev version of SourceTracker via conda:

```
wget https://data.qiime2.org/distro/core/qiime2-2022.2-py38-linux-conda.yml
conda env create -n qiime2-2022.2 --file qiime2-2022.2-py38-linux-conda.yml
rm qiime2-2022.2-py38-linux-conda.yml # cleanup
conda activate qiime2-2022.2
pip install https://github.com/biota/sourcetracker2/archive/master.zip
```

Installation of Qiime1 to access filter_samples_from_otu_table.py

```
conda create -n qiime1 python=2.7 qiime -c bioconda
```

Convert OTU table from *.tsv* to *.biom*.

```
biom convert -i 04-analysis/OTUfilter_table.tsv -o 04-analysis/sourcetracker/OTUfilter-tab
```

Filter OTU table

```
filter_samples_from_otu_table.py \
-i 04-analysis/sourcetracker/OTUfilter-table-from-tsv_json.biom \
-o 04-analysis/sourcetracker/OTUs1000filter_table.biom \
-n 1000
```

Table summary

```
biom summarize-table -i 04-analysis/sourcetracker/OTUs1000filter_table.biom > 04-analysis/
```

Convert to TSV for use with the **decontam** package.

```
biom convert -i 04-analysis/sourcetracker/OTUs1000filter_table.biom -o 04-analysis/deconta
```

Run SourceTracker2 on the filtered OTU table with rarefaction depth of 1000 for both source and samples. Samples and sources were mapped in the ST_comb-plaque-map.txt. The plaque source is a combination of supragingival and subgingival plaque.

```
conda activate qiime2-2022.2
```

```
sourcetracker2 \
    -i 04-analysis/sourcetracker/OTUs1000filter_table.biom \
    -m 04-analysis/sourcetracker/ST_comb-plaque-map.txt \
    --source_sink_column SourceSink  \
    --source_column_value source \
    --source_rarefaction_depth 1000 \
    --sink_rarefaction_depth 1000 \
    --sink_column_value sink \
    --source_category_column Env \
    -o 04-analysis/sourcetracker/sourcetracker2_output \
    --jobs 2 \
    --per_sink_feature_assignments
```

Plots were created of estimated contributions of various sources to the saliva, model calculus and medium samples. Samples are arranged from left to right by how late in the experiment they were sampled, with left being the earliest samples (Figure 1). The output from SourceTracker2 was compared to a dataset of known oral species (Figure 2).

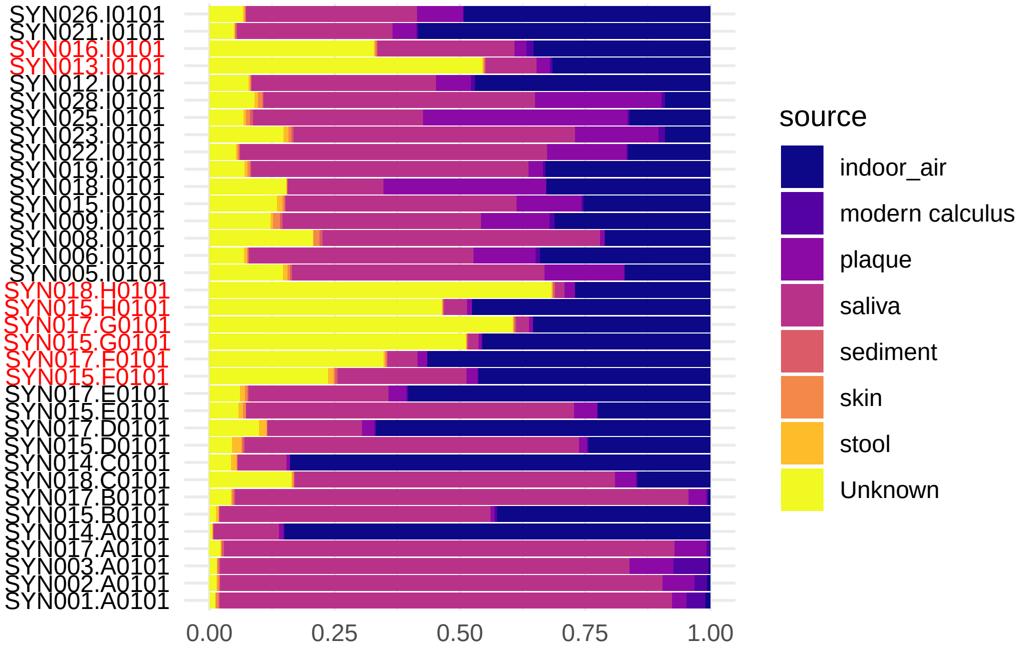

Figure S 1: Estimated proportion of source composition of the abundance-filtered oral biofilm model samples using SourceTracker2. Names of removed samples in red text.

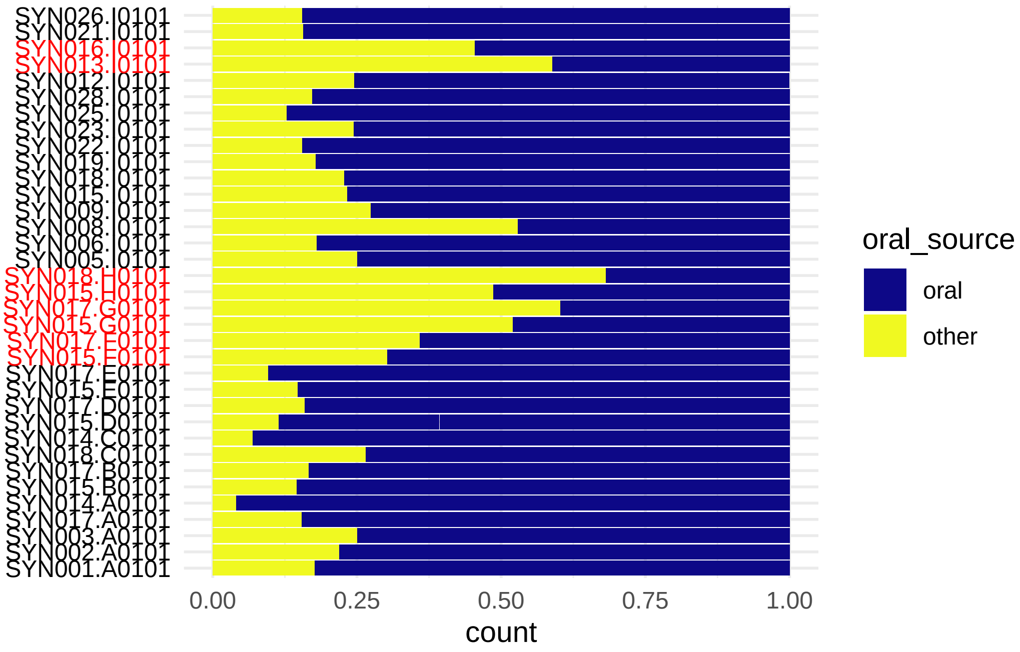

Figure S 2: Estimated proportion of oral or other composition of the abundance-filtered oral biofilm model samples using the oral database from the cuperdec R package. Names of removed samples in red text.

##### decontam

After running `isContaminant()` and filtering out (non-oral) potential contaminants, the samples contained between 88 and 284 species with a mean of 181.59 (Figure 3).

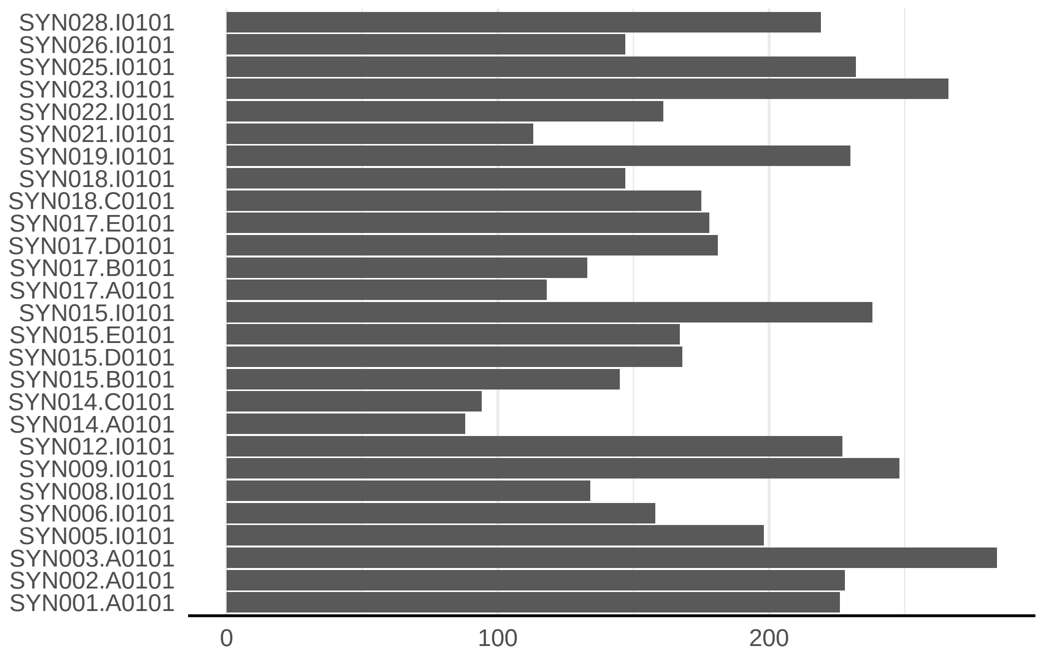

Figure S 3: Species counts after removal of contaminants for biofilm model samples.

### Alpha-diversity

#### Within experiment

There was a slight decrease in mean Shannon Index between inoculation (mean [M] = 2.15 ± 0.808) and treatment samples (M = 1.78 ± 0.568), followed by a slight increase to model calculus samples (M = 1.86 ± 0.261), as well as a decrease in variance within samples types. The Pielou Evenness Index showed a similar pattern (M = 0.41 ± 0.138; M = 0.351 ± 0.108; M = 0.354 ± 0.0379), while number of species increased between the treatment period and the final model calculus (M = 189 ± 82.4; M = 155 ± 29.8; M = 194 ± 49.2).

#### Compared to oral reference samples

We used the Shannon Index to compare alpha-diversity in our model to oral reference samples. The mean Shannon Index of model samples—medium, model calculus, reference *in vitro* biofilm (M = 1.74 ± 0.627; M = 1.86 ± 0.261; M = 1.69 ± 0.173, respectively) were consistently lower than the means of oral reference samples—mucosa, modern reference dental calculus, saliva, and subgingival and subgingival plaque (M = 2.74 ± 0.461; M = 3.14 ± 0.255; M = 3.02 ± 0.548; M = 3.53 ± 0.241; M = 3.04 ± 0.391). The Pielou species evenness index has a similar distribution, although the comparative biofilm samples have a higher mean than biofilm samples from this study. Saliva inoculate samples from this study ($M = 2.5 \pm 0.472$ have a lower mean Shannon index than reference samples ($M = 3.33 \pm 0.297$), which may have contributed to the lower alpha-diversity in model samples compared to reference samples.

### Species composition

Counts from model biofilm and oral reference samples were transformed with a centered ratio log-transform and ordered by PC1 loading (Figure 4 & Figure 5).

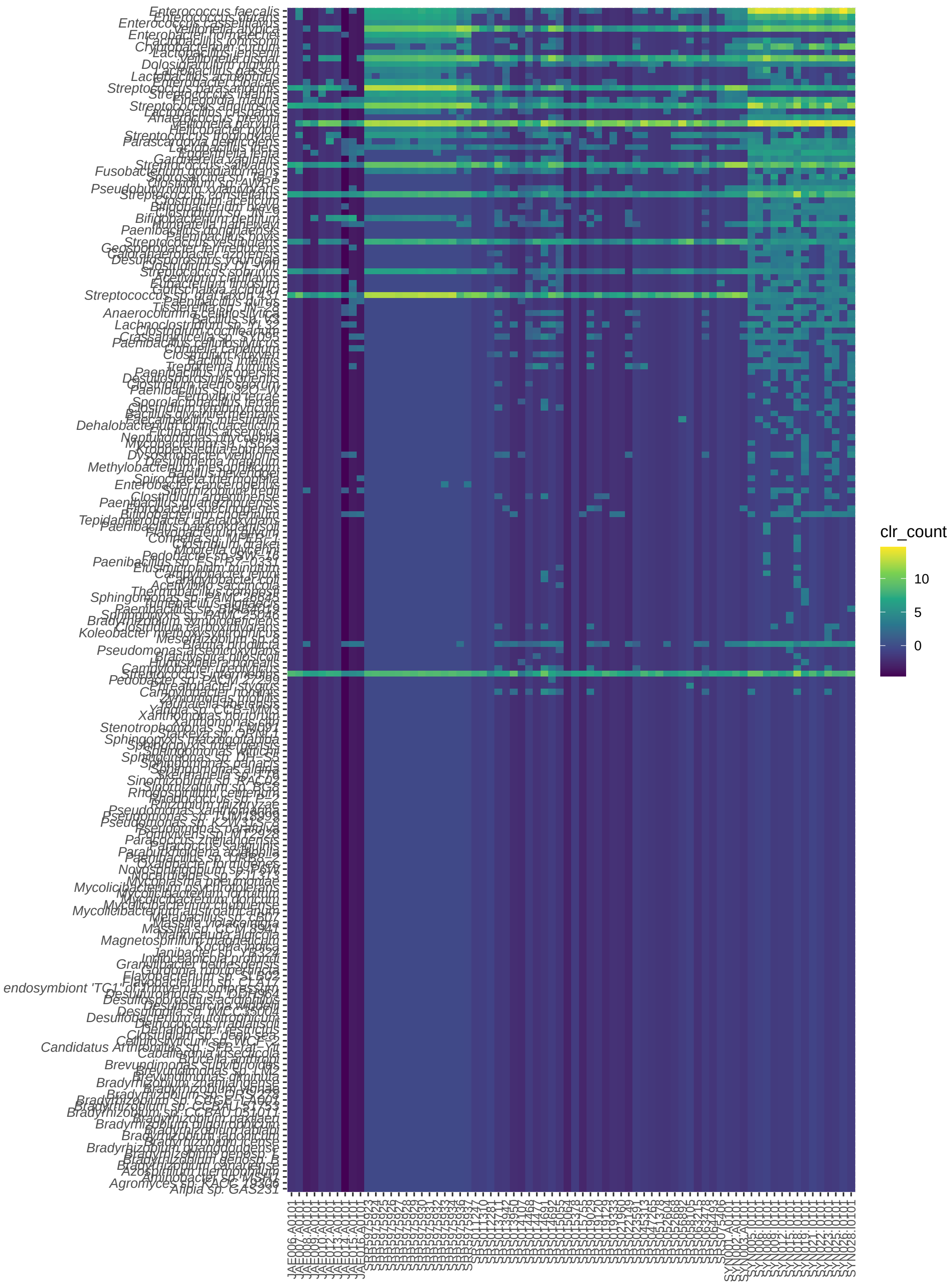

Figure S 4: Centered ratio log-transform abundance of the top 200 species with the highest positive loading on PC1.

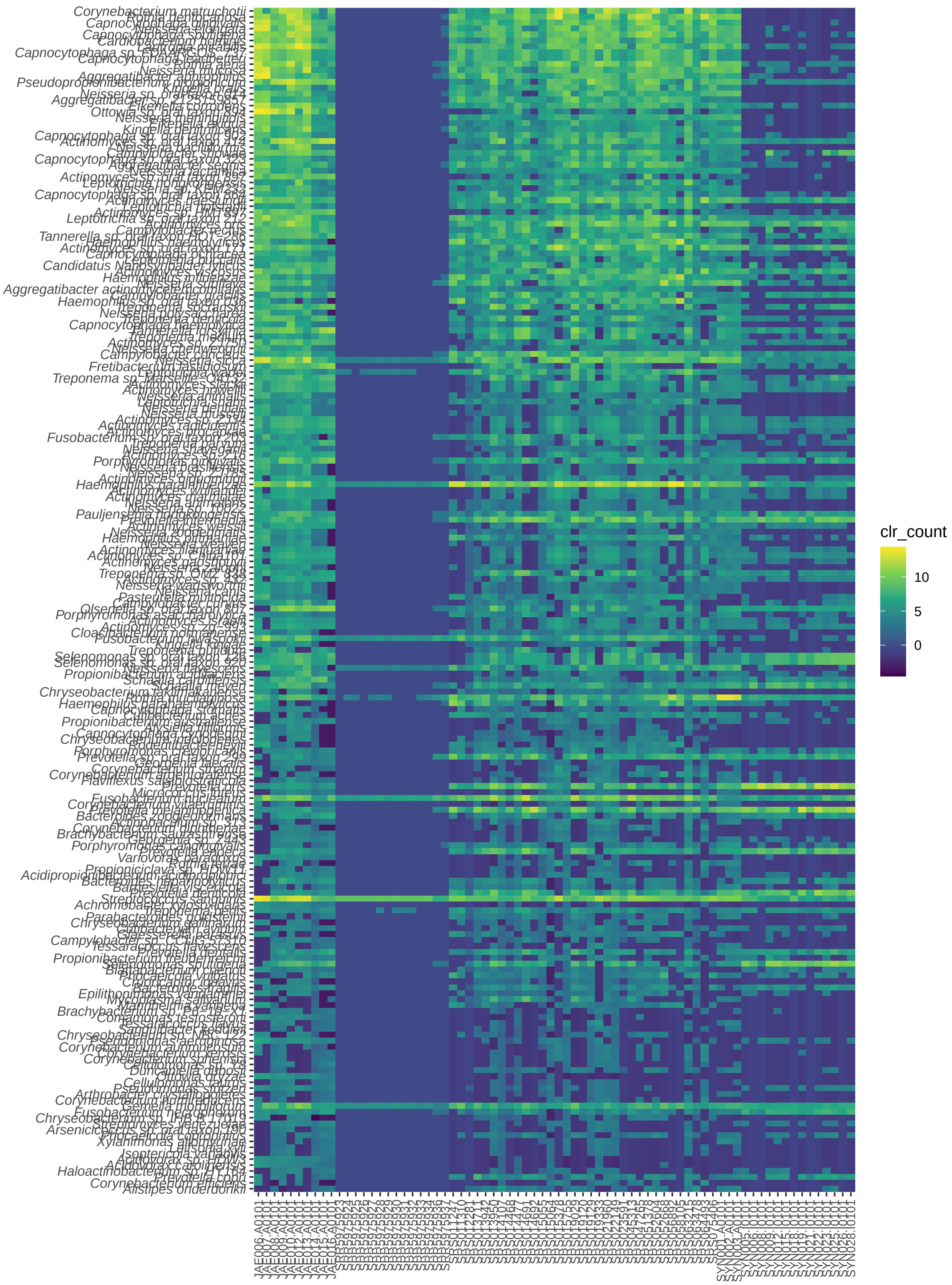

Figure S 5: Centered ratio log-transform abundance of the top 200 species with the highest negative loading on PC1.

### FTIR spectra

Select FTIR spectra not shown in the main manuscript (Figure 6).

Interactive plots are available in the HTML output file.

### Software versions

#### EAGER

Software versions:

| Software | Version |
|---|---|
| nf-core/eager | v2.4.4 |
| Nextflow | v21.03.0.edge |
| FastQC | v0.11.9 |
| MultiQC | v1.12 |
| AdapterRemoval | v2.3.2 |
| fastP | v0.20.1 |
| BWA | v0.7.17-r1188 |
| Bowtie2 | v2.4.4 |
| circulargenerator | v1.0 |
| Samtools | v1.12 |
| endorS.py | v0.4 |
| DeDup | v0.12.8 |
| Picard MarkDuplicates | v2.26.0 |
| Qualimap | v2.2.2-dev |
| Preseq | v3.1.1 |
| GATK HaplotypeCaller | v4.2.0.0 |
| GATK UnifiedGenotyper | v3.5-0-g36282e4 |
| freebayes | v1.3.5 |
| sequenceTools | v1.5.2 |
| VCF2genome | v0.91 |
| MTNucRatioCalculator | v0.7 |
| bedtools | v2.30.0 |
| DamageProfiler | v0.4.9 |
| bamUtil | v1.0.15 |
| pmdtools | v0.50 |
| angsd | v0.935 |
| sexdeterrmine | v1.1.2 |
| multivcfanalyzer | v0.85.2 |
| malt | v0.4.1 |

| Software | Version |
|---|---|
| kraken | v2.1.2 |
| maltextract | v1.7 |
| eigenstrat_snp_coverage | v1.0.2 |
| mapDamage2 | v2.2.1 |
| bbduk | vJanuary 26, 2021 |
| bcftools | v1.12 |

## R

#### Session

```
## R version 4.3.0 (2023-04-21)
## Platform: x86_64-pc-linux-gnu (64-bit)
## Running under: Pop!_OS 22.04 LTS
##
## Matrix products: default
## BLAS:   /usr/lib/x86_64-linux-gnu/blas/libblas.so.3.10.0
## LAPACK: /usr/lib/x86_64-linux-gnu/lapack/liblapack.so.3.10.0
##
## attached base packages:
## [1] stats     graphics  grDevices datasets  utils     methods   base
##
## other attached packages:
## [1] cuperdec_1.1.0 plotly_4.10.1  ggplot2_3.4.2  stringr_1.5.0  readr_2.1.4
## [6] forcats_1.0.0  tidyr_1.3.0    dplyr_1.1.2    here_1.0.1
##
## loaded via a namespace (and not attached):
##  [1] utf8_1.2.3          generics_0.1.3      renv_0.17.3
##  [4] stringi_1.7.12      hms_1.1.3           digest_0.6.31
##  [7] magrittr_2.0.3      evaluate_0.21       grid_4.3.0
## [10] fastmap_1.1.1       rprojroot_2.0.3     jsonlite_1.8.4
## [13] BiocManager_1.30.20 httr_1.4.6          purrr_1.0.1
## [16] fansi_1.0.4         viridisLite_0.4.2   scales_1.2.1
## [19] lazyeval_0.2.2      cli_3.6.1           crayon_1.5.2
## [22] rlang_1.1.1         bit64_4.0.5         munsell_0.5.0
## [25] withr_2.5.0         yaml_2.3.7          parallel_4.3.0
## [28] tools_4.3.0         tzdb_0.3.0          colorspace_2.1-0
## [31] vctrs_0.6.2         R6_2.5.1            lifecycle_1.0.3
## [34] htmlwidgets_1.6.2   bit_4.0.5           vroom_1.6.3
## [37] pkgconfig_2.0.3     pillar_1.9.0        gtable_0.3.3
```

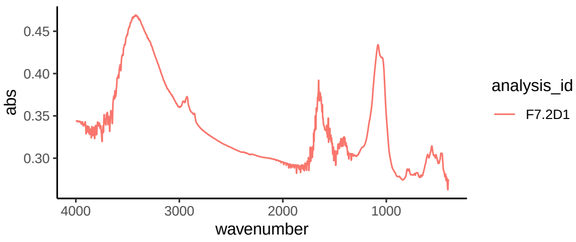

(a) Day 7

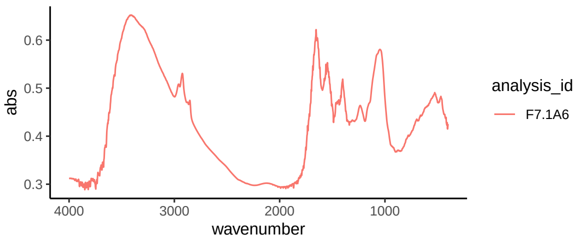

(b) Day 7

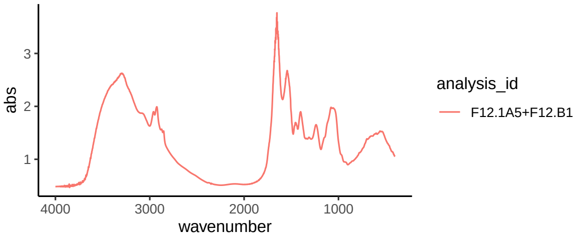

(c) Day 12

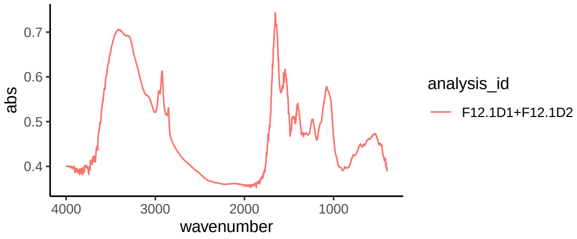

(d) Day 12

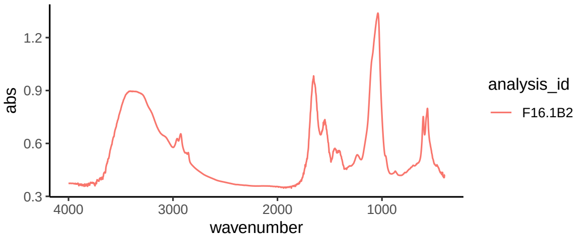

(e) Day 7

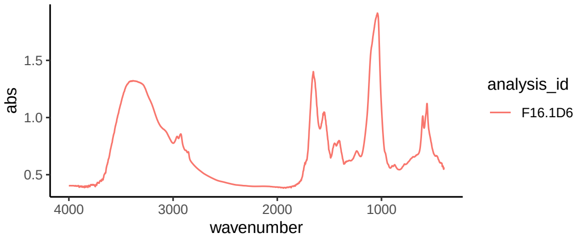

(f) Day 7

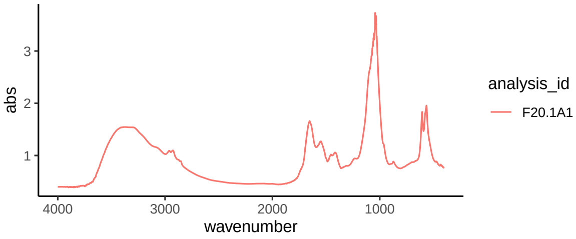

(g) Day 12

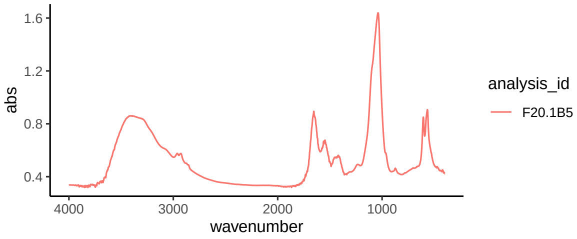

(h) Day 12

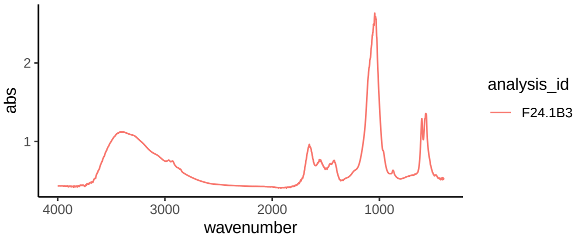

(i) Day 7

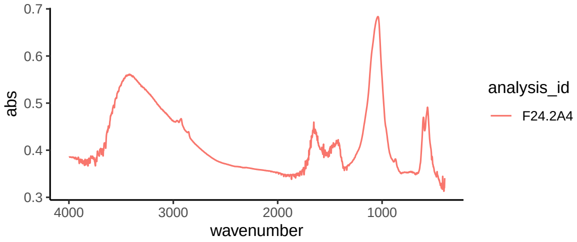

(j) Day 7

Figure S 6: FTIR spectra

```
## [40] glue_1.6.2          data.table_1.14.8   xfun_0.39
## [43] tibble_3.2.1        tidyselect_1.2.0    rstudioapi_0.14
## [46] knitr_1.42          farver_2.1.1        htmltools_0.5.5
## [49] labeling_0.4.2      rmarkdown_2.21      compiler_4.3.0
```

#### Packages

Table 6: List of R packages and dependencies.

| Package | Version | Source |
|---|---|---|
| ANCOMBC | 2.2.0 | Bioconductor |
| BH | 1.81.0-1 | Repository |
| Biobase | 2.60.0 | Bioconductor |
| BiocBaseUtils | 1.2.0 | Bioconductor |
| BiocGenerics | 0.46.0 | Bioconductor |
| BiocManager | 1.30.20 | Repository |
| BiocNeighbors | 1.18.0 | Bioconductor |
| BiocParallel | 1.34.1 | Bioconductor |
| BiocSingular | 1.16.0 | Bioconductor |
| Biostrings | 2.68.0 | Bioconductor |
| CVXR | 1.0-11 | Repository |
| Cairo | 1.6-0 | Repository |
| DBI | 1.1.3 | Repository |
| DECIPHER | 2.28.0 | Bioconductor |
| DEoptimR | 1.0-13 | Repository |
| DelayedArray | 0.26.2 | Bioconductor |
| DelayedMatrixStats | 1.22.0 | Bioconductor |
| DescTools | 0.99.48 | Repository |
| DirichletMultinomial | 1.42.0 | Bioconductor |
| ECOSolveR | 0.5.4 | Repository |
| Exact | 3.2 | Repository |
| FNN | 1.1.3.2 | Repository |
| Formula | 1.2-5 | Repository |
| GenomeInfoDb | 1.36.0 | Bioconductor |
| GenomeInfoDbData | 1.2.10 | Bioconductor |
| GenomicRanges | 1.52.0 | Bioconductor |
| Hmisc | 5.1-0 | Repository |
| IRanges | 2.34.0 | Bioconductor |
| MASS | 7.3-59 | Repository |
| Matrix | 1.5-1 | Repository |
| MatrixGenerics | 1.12.0 | Bioconductor |
| MultiAssayExperiment | 1.26.0 | Bioconductor |

| Package | Version | Source |
|---|---|---|
| R6 | 2.5.1 | Repository |
| RColorBrewer | 1.1-3 | Repository |
| RCurl | 1.98-1.12 | Repository |
| RSQLite | 2.3.1 | Repository |
| RSpectra | 0.16-1 | Repository |
| Rcpp | 1.0.10 | Repository |
| RcppAnnoy | 0.0.20 | Repository |
| RcppArmadillo | 0.12.2.0.0 | Repository |
| RcppEigen | 0.3.3.9.3 | Repository |
| RcppHNSW | 0.4.1 | Repository |
| RcppML | 0.3.7 | Repository |
| RcppProgress | 0.4.2 | Repository |
| RcppTOML | 0.2.2 | Repository |
| Rdpack | 2.4 | Repository |
| Rhdf5lib | 1.22.0 | Bioconductor |
| Rmpfr | 0.9-2 | Repository |
| Rtsne | 0.16 | Repository |
| S4Arrays | 1.0.1 | Bioconductor |
| S4Vectors | 0.38.1 | Bioconductor |
| ScaledMatrix | 1.8.1 | Bioconductor |
| SingleCellExperiment | 1.22.0 | Bioconductor |
| SummarizedExperiment | 1.30.1 | Bioconductor |
| TreeSummarizedExperiment | 2.8.0 | Bioconductor |
| XVector | 0.40.0 | Bioconductor |
| ade4 | 1.7-22 | Repository |
| ape | 5.7-1 | Repository |
| askpass | 1.1 | Repository |
| assertthat | 0.2.1 | Repository |
| backports | 1.4.1 | Repository |
| base64enc | 0.1-3 | Repository |
| basilisk | 1.12.0 | Bioconductor |
| basilisk.utils | 1.12.0 | Bioconductor |
| bayesm | 3.1-5 | Repository |
| beachmat | 2.16.0 | Bioconductor |
| beeswarm | 0.4.0 | Repository |
| biomformat | 1.28.0 | Bioconductor |
| bit | 4.0.5 | Repository |
| bit64 | 4.0.5 | Repository |
| bitops | 1.0-7 | Repository |
| blob | 1.2.4 | Repository |
| boot | 1.3-28 | Repository |

| Package | Version | Source |
| --- | --- | --- |
| brio | 1.1.3 | Repository |
| bslib | 0.4.2 | Repository |
| cachem | 1.0.8 | Repository |
| callr | 3.7.3 | Repository |
| cellranger | 1.1.0 | Repository |
| checkmate | 2.2.0 | Repository |
| class | 7.3-21 | Repository |
| cli | 3.6.1 | Repository |
| clipr | 0.8.0 | Repository |
| cluster | 2.1.4 | Repository |
| codetools | 0.2-19 | Repository |
| colorspace | 2.1-0 | Repository |
| compositions | 2.0-6 | Repository |
| corpcor | 1.6.10 | Repository |
| cowplot | 1.1.1 | Repository |
| cpp11 | 0.4.3 | Repository |
| crayon | 1.5.2 | Repository |
| crosstalk | 1.2.0 | Repository |
| cuperdec | 1.1.0 | Repository |
| curl | 5.0.0 | Repository |
| data.table | 1.14.8 | Repository |
| decontam | 1.20.0 | Bioconductor |
| densvis | 1.10.1 | Bioconductor |
| desc | 1.4.2 | Repository |
| diffobj | 0.3.5 | Repository |
| digest | 0.6.31 | Repository |
| dir.expiry | 1.8.0 | Bioconductor |
| doParallel | 1.0.17 | Repository |
| doRNG | 1.8.6 | Repository |
| dplyr | 1.1.2 | Repository |
| dqrng | 0.3.0 | Repository |
| e1071 | 1.7-13 | Repository |
| ellipse | 0.4.5 | Repository |
| ellipsis | 0.3.2 | Repository |
| emmeans | 1.8.5 | Repository |
| energy | 1.7-11 | Repository |
| estimability | 1.4.1 | Repository |
| evaluate | 0.21 | Repository |
| expm | 0.999-7 | Repository |
| fansi | 1.0.4 | Repository |
| farver | 2.1.1 | Repository |

| Package | Version | Source |
|---|---|---|
| fastmap | 1.1.1 | Repository |
| filelock | 1.0.2 | Repository |
| fontawesome | 0.5.1 | Repository |
| forcats | 1.0.0 | Repository |
| foreach | 1.5.2 | Repository |
| foreign | 0.8-82 | Repository |
| formatR | 1.14 | Repository |
| fs | 1.6.2 | Repository |
| futile.logger | 1.4.3 | Repository |
| futile.options | 1.0.1 | Repository |
| generics | 0.1.3 | Repository |
| ggbeeswarm | 0.7.2 | Repository |
| ggplot2 | 3.4.2 | Repository |
| ggrastr | 1.0.1 | Repository |
| ggrepel | 0.9.3 | Repository |
| gld | 2.6.6 | Repository |
| glue | 1.6.2 | Repository |
| gmp | 0.7-1 | Repository |
| gridExtra | 2.3 | Repository |
| gsl | 2.1-8 | Repository |
| gtable | 0.3.3 | Repository |
| here | 1.0.1 | Repository |
| highr | 0.10 | Repository |
| hms | 1.1.3 | Repository |
| htmlTable | 2.4.1 | Repository |
| htmltools | 0.5.5 | Repository |
| htmlwidgets | 1.6.2 | Repository |
| httr | 1.4.6 | Repository |
| igraph | 1.4.2 | Repository |
| irlba | 2.3.5.1 | Repository |
| isoband | 0.2.7 | Repository |
| iterators | 1.0.14 | Repository |
| jquerylib | 0.1.4 | Repository |
| jsonlite | 1.8.4 | Repository |
| knitr | 1.42 | Repository |
| labeling | 0.4.2 | Repository |
| lambda.r | 1.2.4 | Repository |
| later | 1.3.1 | Repository |
| lattice | 0.21-8 | Repository |
| lazyeval | 0.2.2 | Repository |
| lifecycle | 1.0.3 | Repository |

| Package | Version | Source |
|---|---|---|
| lme4 | 1.1-33 | Repository |
| lmerTest | 3.1-3 | Repository |
| lmom | 2.9 | Repository |
| magrittr | 2.0.3 | Repository |
| matrixStats | 0.63.0 | Repository |
| memoise | 2.0.1 | Repository |
| mgcv | 1.8-42 | Repository |
| mia | 1.8.0 | Bioconductor |
| microbiome | 1.22.0 | Bioconductor |
| mime | 0.12 | Repository |
| minqa | 1.2.5 | Repository |
| mixOmics | 6.24.0 | Repository |
| multtest | 2.56.0 | Bioconductor |
| munsell | 0.5.0 | Repository |
| mvtnorm | 1.1-3 | Repository |
| nlme | 3.1-162 | Repository |
| nloptr | 2.0.3 | Repository |
| nnet | 7.3-18 | Repository |
| numDeriv | 2016.8-1.1 | Repository |
| openssl | 2.0.6 | Repository |
| osqp | 0.6.0.8 | Repository |
| patchwork | 1.1.2 | Repository |
| permute | 0.9-7 | Repository |
| pheatmap | 1.0.12 | Repository |
| phyloseq | 1.44.0 | Bioconductor |
| pillar | 1.9.0 | Repository |
| pixmap | 0.4-12 | Repository |
| pkgconfig | 2.0.3 | Repository |
| pkgload | 1.3.2 | Repository |
| plogr | 0.2.0 | Repository |
| plotly | 4.10.1 | Repository |
| plyr | 1.8.8 | Repository |
| png | 0.1-8 | Repository |
| praise | 1.0.0 | Repository |
| prettyunits | 1.1.1 | Repository |
| processx | 3.8.1 | Repository |
| progress | 1.2.2 | Repository |
| promises | 1.2.0.1 | Repository |
| proxy | 0.4-27 | Repository |
| ps | 1.7.5 | Repository |
| purrr | 1.0.1 | Repository |

| Package | Version | Source |
|---|---|---|
| rARPACK | 0.11-0 | Repository |
| ragg | 1.2.5 | Repository |
| rappdirs | 0.3.3 | Repository |
| rbbt | 0.0.0.9000 | GitHub |
| rbibutils | 2.2.13 | Repository |
| readr | 2.1.4 | Repository |
| readxl | 1.4.2 | Repository |
| rematch | 1.0.1 | Repository |
| rematch2 | 2.1.2 | Repository |
| renv | 0.17.3 | Repository |
| reshape2 | 1.4.4 | Repository |
| reticulate | 1.28 | Repository |
| rhdf5 | 2.44.0 | Bioconductor |
| rhdf5filters | 1.12.1 | Bioconductor |
| rlang | 1.1.1 | Repository |
| rmarkdown | 2.21 | Repository |
| rngtools | 1.5.2 | Repository |
| robustbase | 0.95-1 | Repository |
| rootSolve | 1.8.2.3 | Repository |
| rpart | 4.1.19 | Repository |
| rprojroot | 2.0.3 | Repository |
| rstudioapi | 0.14 | Repository |
| rsvd | 1.0.5 | Repository |
| sass | 0.4.6 | Repository |
| scales | 1.2.1 | Repository |
| scater | 1.28.0 | Bioconductor |
| scs | 3.2.4 | Repository |
| scuttle | 1.10.1 | Bioconductor |
| sitmo | 2.0.2 | Repository |
| snow | 0.4-4 | Repository |
| sp | 1.6-0 | Repository |
| sparseMatrixStats | 1.12.0 | Bioconductor |
| stringi | 1.7.12 | Repository |
| stringr | 1.5.0 | Repository |
| survival | 3.5-3 | Repository |
| sys | 3.4.1 | Repository |
| systemfonts | 1.0.4 | Repository |
| tensorA | 0.36.2 | Repository |
| testthat | 3.1.8 | Repository |
| textshaping | 0.3.6 | Repository |
| tibble | 3.2.1 | Repository |

| Package | Version | Source |
|---|---|---|
| tidyr | 1.3.0 | Repository |
| tidyselect | 1.2.0 | Repository |
| tidytree | 0.4.2 | Repository |
| tinytex | 0.45 | Repository |
| treeio | 1.24.0 | Bioconductor |
| tzdb | 0.3.0 | Repository |
| utf8 | 1.2.3 | Repository |
| uwot | 0.1.14 | Repository |
| vctrs | 0.6.2 | Repository |
| vegan | 2.6-4 | Repository |
| vipor | 0.4.5 | Repository |
| viridis | 0.6.3 | Repository |
| viridisLite | 0.4.2 | Repository |
| vroom | 1.6.3 | Repository |
| waldo | 0.5.1 | Repository |
| withr | 2.5.0 | Repository |
| xfun | 0.39 | Repository |
| yaml | 2.3.7 | Repository |
| yulab.utils | 0.0.6 | Repository |
| zlibbioc | 1.46.0 | Bioconductor |
